# Increased pathogenicity of the nematophagous fungus *Drechmeria coniospora* following long-term laboratory culture

**DOI:** 10.1101/2021.09.01.457749

**Authors:** Damien Courtine, Xing Zhang, Jonathan J. Ewbank

**Affiliations:** Aix Marseille Univ, CNRS, INSERM, CIML, Turing Centre for Living Systems, Marseille, France

## Abstract

Domestication provides a window into adaptive change. Over the course of 2 decades of laboratory culture, a strain of the nematode-specific fungus *Drechmeria coniospora* became more virulent during its infection of *Caenorhabditis elegans*. Through a close comparative examination of the genome sequences of the original strain and its more pathogenic derivative, we identified a small number of non-synonymous mutations in protein-coding genes. In one case, the mutation was predicted to affect a gene involved in hypoxia resistance and we provide direct corroborative evidence for such an effect. The mutated genes with functional annotation were all predicted to impact the general physiology of the fungus and this was reflected in an increased *in vitro* growth, even in the absence of *C. elegans*. While most cases involved single nucleotide substitutions predicted to lead to a loss of function, we also observed a predicted restoration of gene function through deletion of an extraneous tandem repeat. This latter change affected the regulatory subunit of a cAMP-dependent protein kinase. Remarkably, we also found a mutation in a gene for a second protein of the same, protein kinase A, pathway. Together, we predict that they result in a stronger repression of the pathway for given levels of ATP and adenylate cyclase activity. Finally, we also identified mutations in a few lineage-specific genes of unknown function that are candidates for factors that influence virulence in a more direct manner.

## Introduction

*Drechmeria coniospora* is a nematophagous *Ascomycetes* fungus that lies on a distal branch of the *Ophiocordycipitaceae* (Li et al., 2021). It is one of the best-characterised fungal pathogens of *Caenorhabditis elegans*. Infection starts with the adhesion of non-motile asexual spores (conidia) to the nematode cuticle. These single-celled haploid spores are formed through a phialidic mode of holoblastic conidiogenesis. In other words, they grow out from a conidiogenic hypha, on specialized stalks called conidiophores, with their extension involving the complete cell wall of the hypha, and can be distinguished from the conidiogenic hypha before they separate from it. They are also called mitospores, as they are generated through mitosis, with no meiosis, and are therefore genetically identical to their haploid parent. After adhesion, an appressorium forms, allowing the nematode cuticle to be penetrated. Haploid and septate endozoic hyphae grow throughout the infected host, with new conidiophores then emerging through the cuticle of the nematode, forming conidia that can go on to infect other nematodes (Gernandt and Stone, 1999; Saikawa, 1982; Wyatt et al., 2013).

A variety of isolates from diverse locations worldwide exist, including ATCC 96282, derived from a strain collected in Sweden by H.-B. Jansson in 1990 (Jansson and Friman, 1999). This strain has been extensively used in studies addressing host defences (Dierking et al., 2011; Labed et al., 2012; Lee et al., 2010, 2018). More recently, taking its genome as a starting point, *D. coniospora* has been developed as a model to understand how nematophagous fungi can infect and kill their hosts. This capacity has emerged multiple times during evolution and in each case appears largely to involve distinct molecular mechanisms (Andersson et al., 2014; Ji et al., 2020; Lebrigand et al., 2016; Lin et al., 2018; Meerupati et al., 2013; Wang et al., 2018; Xie et al., 2016), although there is also evidence in some cases for convergent evolution (Iqbal et al., 2018).

With regards *D. coniospora*, as a first example, presumably as a consequence of horizontal gene transfer, the fungus has acquired genes encoding proteins containing one or more SapA domains that bind to and potentially inhibit the antimicrobial activity of host SapB-domains proteins (Lebrigand et al., 2016). Secondly, *D. coniospora* has an unusually broad repertoire of enterotoxins (Wang et al., 2018). Only two out of these 23 enterotoxins have been characterised in any detail. They play specific, partly antagonistic roles, interfering with host defence signalling pathways (Zhang et al., 2021). These examples represent a tiny fraction of the hundreds of potential virulence factors encoded within the *D. coniospora* genome. A substantial proportion of them are either lineage-specific or correspond to proteins for which no functional annotation exists in any species (Lebrigand et al., 2016). Many aspects of the biology of *D. coniospora* that are most relevant to its capacity to parasitise worms are therefore poorly characterised.

One powerful way to address the molecular basis of physiological traits is to take advantage of genetic and phenotypic variation between natural populations. The genome of a Danish isolate of *D. coniospora* has also been sequenced (Zhang et al., 2016), but its genome is very different from that of ATCC 96282 (Courtine et al., 2020), rendering comparative functional analysis challenging. Another approach relies on experimental evolution. Typically, organisms are maintained in a defined environment, with or without applied selective pressures, and the changes that accumulate over time are monitored. The best known such study is one from the Lenski lab, where 12 populations of *Escherichia coli* have been cultured since 1988 in minimal media by batch culture for more than 60,000 generations (Good et al., 2017; Lenski, 2017). In another long-running experiment, *Saccharomyces cerevisiae* populations evolved over 3 years, for some 10,000 generations in three environments (Johnson et al., 2021; McDonald, 2019). These and other studies have given valuable insights into evolutionary processes. It has been observed repeatedly that populations follow a more-or-less foreseeable path, with the rate that fitness increases slowing down as populations adapt, while the mutation rate remains relatively stable. In comparison to complex and changing natural environments, laboratory culture conditions are designed to be simple and constant. Although it is impossible to predict what exact mutations will arise, selection pressure will generally favour the loss of genes and pathways that are superfluous under the experimental conditions (Kvitek and Sherlock, 2013; McDonald, 2019).

Experimental evolution has also been used to characterise host-pathogen interactions. In the case of *C. elegans*, pioneering studies include a dissection of the role of mating modalities in host resistance (Morran et al., 2011), behavioural responsiveness to bacterial microbes (Schulte et al., 2012), and co-evolutionary studies (Masri et al., 2015). The field continues to be very active (e.g. (Ekroth et al., 2021; White et al., 2020, 2021)). Here, we took advantage of an unintentional domestication experiment. In 1999, we started laboratory culture of the *D. coniospora* ATCC 96282 strain. We recently sequenced the genome of this isolate, which we refer to as Swe1, as well as that of a derived strain, Swe3, cryo-archived in 2018 after almost 2 decades of laboratory culture (Courtine et al., 2020). As we show here, there is a notable difference in the speed at which Swe1 and Swe3 kill *C. elegans*.

Leveraging the genome annotation available for the Swe1 derivative Swe2, cryo-archived and sequenced in 2013 (Lebrigand et al., 2016), we were able to annotate the Swe1 and Swe3 genomes. This then provided an opportunity to conduct a comparative genomic analysis, to identify the mutations that have accumulated over the years, including those potentially linked to the observed increase in virulence. When the laboratory culture of *D. coniospora* was started, there was little expectation that it would continue for so long. Had this been the case, a more controlled experimental protocol would have been instigated, with, for example, passaging of spores at more regular intervals, and replicate populations grown in parallel to be able to determine the robustness and reproducibility of any observed changes. Nevertheless, this single somewhat uncontrolled long-term experiment does provide insight into the evolutionary changes that occurred during domestication of *D. coniospora* and that impact its virulence

## Methods

### *C*.*elegans* strains and culture

The strain IG463 (*rrf-3(b26);frIs7*) was made by standard crosses between the conditionally sterile mutant strain DH26 *rrf-3(b26) II* (formerly *fer-15*) and the reporter strain IG274 (+;*frIs7* [*nlp-29p::GFP, col-12p::DsRed*] *IV* (Pujol et al., 2008). N2 and other strains were maintained on nematode growth media (NGM) and fed *E. coli* strain OP50 (Stiernagle, 2006). IG463 and DH26 were maintained at 15°C, the permissive temperature, the other strains at 25°C.

### *D. coniospora* culture

*D. coniospora* was serially cultured by infecting *C. elegans*. Typically, spores were harvested from infected worms every one or two weeks and used to infect a fresh worm population. The methods used are described in detail elsewhere (Powell and Ausubel, 2008). Briefly, about 300 μl of freshly harvested spore solution (ca. 1-5 × 10^8^ spores) was added to a standard 10 cm NGM agar plate with 1000-2000 synchronized L4 or young adult N2 worms on an extensive lawn of *E. coli* OP50. After drying under a laminar flow hood, the plate was incubated at 25°C for 1 day. Infected worms were harvested with 50 mM NaCl and transferred to an NGM plate supplemented with 15 μg/ml gentamicin and 100 μg/ml ampicillin, without OP50. The plate was incubated at 25°C for up to 1 week and then stored at 20°C.

### Infection assays

Spores were harvested from plates at 25°C after 6 days and counted using a Neubauer chamber (Bürker) as described (http://www.lo-laboroptik.de/englisch/info/info.html) before dilution in 50 mM NaCl to the required concentration. Around 150 young adult IG463 worms that had been grown at 25°C, the non-permissive temperature, were manually transferred to 4 cm plates containing NGM agar seeded with *E. coli* OP50. Freshly harvested spores (1 × 10^9^) were spread on the plate. After drying under a laminar flow hood, and overnight infection at 25°C, for each experimental condition, 25 worms were picked into 4 wells of a 12-well plate containing NGM agar seeded with *E. coli* OP50. Images of each well were collected automatically every 24 minutes using a custom system that will be described elsewhere. The images were examined, and worms scored as dead when they no longer exhibited movement between successive images.

### Analyses with the Biosort worm sorter

Fluorescent protein expression was quantified with the COPAS (Complex Object Parametric Analyzer and Sorter) Biosort system (Union Biometrica, Holliston, MA) as described (Pujol et al., 2008). Worms were analyzed for length (assessed as TOF, time of flight), optical density (assessed as extinction) and Green and/or Red fluorescence (GFP/Red). Raw data were filtered on the TOF for adult worms (typically 300 ≤ TOF ≤ 1500). Statistical significance was determined using an unpaired t-test (GraphPad Prism).

### Monitoring fungal growth

Spores (typically 10^6^ in 30 µl) were added to each well of 12-well plates containing 700 µl of NGM agar supplemented with 0 mM, 1 mM, or 2 mM CoCl_2_ and incubated at 25°C. Fungal growth was recorded using an Zeiss Axio Observer Z1 microscope equipped with Definite Focus, a motorised stage, a PeCon GmbH incubation chamber and a Hamamatsu C11440-42U30 camera. All images from a stack were aligned using the ImageJ macro *Align_Slice* (https://github.com/landinig/IJ-Align_Slice/) to compensate for any image drift. Spore and hyphae length were measured using the ImageJ toolbox HyphaTracker v1.0 (Brunk et al., 2018; Schindelin et al., 2012). The thresholds to convert images into binary data were set automatically, the minimal area was set to 5 pixels and the last frame was selected to be the reference.

### Fungal PCR

Fungal DNA was extracted as previously described (Courtine et al., 2020) from freshly thawed aliquots of the original isolate ATCC 96282, called here Swe1, or its derivatives Swe2 and Swe3, archived and sequenced in 2013 (Lebrigand et al., 2016) and 2018 (Courtine et al., 2020), respectively. Once samples of Swe1, Swe2 or Swe3, which were cryopreserved at -80°C, were thawed, their precise culture history before being used in experiments was recorded (number of passages, etc). The identity of the different strains was verified regularly by PCR using primers designed to amplify one locus per chromosome, divergent either between Swe1 and Swe2, or between Swe2 and Swe3. The strains could be distinguished on the basis of the results of PCR amplification: presence or absence of an amplicon, size of an amplicon, or amplicon sequence, depending on the strain and PCR primers used (Supp. Table S1). PCR with the indicated primers pairs was also used to produce amplicons from Swe2 genomic DNA, corresponding to regions that were previously poorly defined, that were then sequenced.

### Assembling the Swe2 mitochondrial genome

The genomic reads from Swe2 (SRR1810847) mapping on the Swe1 and Swe3 mitochondrial sequences (Courtine et al., 2020) were extracted and used for a *de novo* assembly with IDBA-UD v1.1.3-1 using default parameters (Peng et al., 2012). The resulting longest contig was circularised with nucmer (*--maxmatch --nosimplify*), followed by show-coords (*-lrcT*) within the package MUMmer v4.0.0b2 (Marçais et al., 2018). This generated a mitochondrial genome that was judged complete on the basis of comparison with available mitochondrial genomic sequences from the closely-related fungi *Purpureocillium lilacinum, Tolypocladium inflatum* and *Tolypocladium cylindrosporum* (Supp. Table S2; Supp. Fig. S1).

### Scaffolding of Swe2 sequences into chromosomal assemblies

A previous comparison of Swe1 and Swe3 genomes revealed a chromosomal collinearity. This had allowed the major Swe2 scaffolds to be assembled into 3 chromosomes (Courtine et al., 2020), but omitted smaller scaffolds that included some predicted protein-coding genes (Supp. Table S3). Each of these unincorporated scaffolds was compared to the Swe1 and Swe3 sequences using BLASTN (Altschul et al., 1997). The alignments were visualised using Kablammo (Wintersinger and Wasmuth, 2015), permitting manual assignment of scaffolds containing repeat sequences to unique chromosomal positions, as well as trimming of unaligned regions. The precise insertion position and orientation for each scaffold was determined by manual inspection of a multiple sequence alignment from the 3 genomes using Mauve (Darling et al., 2004). When the precise junctional sequence could not be defined, an *NNN* sequence was added at either side of each newly inserted scaffold.

### Improving the Swe2 genome sequence

The reassembled Swe2 genome still included 1,386 regions of sequence undetermined in the original assembly *i*.*e*. stretches with one or several consecutives *N*s. Sequences 1000 nt 5’ and 3’ to each of these regions were extracted separately from the Swe2 genome and aligned by BLASTN against the Swe1 and Swe3 genomes. In the simplest case, when unambiguous syntenic alignments revealed regions of defined genome sequence fully conserved between Swe1 and Swe3, the relevant sequence was copied from Swe1/Swe3 and inserted in the place of the undefined region of the Swe2 genome. In cases where the Swe1 and Swe3 sequences had the same length but were not identical, the differences were examined, and only the ones with 10 or less ambiguous positions were retained. These remaining undefined bases were left as *N* before using the consensus sequence to edit the Swe2 genome. This resulted in an addition of 41,997 bp of defined sequence. No attempt was made to correct the Swe2 sequence when the corresponding regions in the Swe1 and Swe3 genomes were of different lengths. In some cases, the flanking regions overlapped when aligned, because of an incorrect copy number for a repetitive sequence in the Swe2 genome. We then used the consensus Swe1 and Swe3 sequences to remove this superfluous sequence from the Swe2 genome, representing a total of 7,579 bp of deleted sequence. As a complementary automatic approach, in parallel we used the tool Sealer, from the assembler ABySS (Paulino et al., 2015) with the parameters “*-b20G -k120 -k110 -k100 -k90 -k80 -k70 -k60 -k50 -k40 -F 700 -P 10 -B 3000*” to generate a list of gaps to be closed, many of which were also identified by the manual workflow. We filtered these out, as well as those containing ambiguous bases, and then implemented the remaining modifications identified by Sealer. The improved version of Swe2 still contains 33,795 *N*s in 993 stretches.

### Genome similarity metrics

Average Nucleotide Identity values were calculated using pyani (v0.2.9) with default parameters (Pritchard et al., 2015). Genomes were aligned with Minimap v2.17-r974-dirty with the parameters *-c --cs=long* (Li, 2018a). The resulting PAF file was converted to MAF with the Minimap2 utility *paftools*.*js view -f maf* and the command *sed ‘s/^a /a score=/*’. The gap-compressed identity score was extracted from the field *de:f* in the PAF file, and the gap-excluded identity score was obtained by parsing the MAF alignment (BioPython AlignIO v1.78). For all gap-free positions, the number of matches and mismatches was recorded, allowing the number of matches divided by number of matches and mismatches to be calculated. These metrics were calculated for alignment blocks longer than 1 Mb (Li, 2018b). Homopolymers were removed from the genomes with the command “sed -e ‘s/[aA]\{5,\}//g’ -e ‘s/[tT]\{5,\}//g’ -e ‘s/[cC]\{5,\}//g’ -e ‘s/[gG]\{5,\}//g’ “.

### Swe2 gene set

A number of proteins predicted for *D. coniospora* ARSEF 6962 (Zhang et al., 2016), referred to here as Dan2, were absent from the original Swe2 protein set as judged by the results of reciprocal BLASTP searches. These included members of highly repeated families, for example integrases and transposases, as well as a few with atypical structures (KYK54054.1/KYK54053.1/KYK56388.1/KYK54065.1)) that we did not attempt to curate. The remaining Dan2 protein sequences were aligned by TBLASTN against the Swe2 genome. Those matching an existing predicted Swe2 protein-coding gene were considered likely to reflect events of gene expansion in Dan2 and not pursued further, with the exception of genes encoding proteins with an enterotoxin alpha domain (PFAM: PF01375), which was the subject of manual annotation (Zhang et al., 2021). Then Dan2 protein sequences that aligned with predicted coding sequence from Swe2 with an identity greater than 90% and for which the main exon was more than 80% of the predicted coding sequence for the gene were manually curated and added to the set of predicted protein-coding genes in Swe2. When polishing of the Swe2 genome removed one or more *N* from a predicted protein-coding gene, the RNAseq reads (SRR1930119, SRR1930124) and/or BLASTX alignment at NCBI against nr/nt databases were used to validate the corrected sequence.

### Gene Comparison

To identify the mutations that accumulated within protein-coding genes, the Swe2 gene set was directly compared to the Swe1 and Swe3 genome sequences, taking advantage of their conserved genomic structure. Thus, from the start of each chromosome, Swe2 genes were sequentially aligned on Swe1 and Swe3 by BLASTN v2.8.1+ with the parameters *-max_hsps 1 -dust no*, verifying at each iteration that the target gene was in the expected chromosomal position. For each such gene, the query coverage, percent of identity, as well as alignment start and stop coordinates on the Swe1 and Swe3 chromosomes were recorded.

We then selected, i) genes with a query coverage equal to 1 and less than 100% sequence identity, ii) genes with a query coverage different from 1. For the former set, exons for each gene were aligned to the expected gene sequence in Swe1 and Swe3 by BLASTN to identify genes with mutations in introns; these were excluded from further analysis. The protein sequence of the remaining genes was aligned to Swe1 and Swe3 by TBLASTN with parameters *-seq no* and *-subject_loc* to define the gene coordinates on the target genome. All high scoring pairs (HSPs) were compared 2 by 2 between Swe1 and Swe3 to identify potential sequence changes. The predicted protein sequence of candidate genes with expected non-synonymous substitutions were aligned on the expected Swe1/Swe3 gene using Exonerate v2.2.0 (Slater and Birney, 2005) and parameters *-m protein2genome* to confirm the mutation. Finally, the sequence evidence for each potentially mutated position was checked in the 3 genomes using the sets of short-reads available: ERR3997395, SRR1810847 and ERR3997392 for Swe1, Swe2 and Swe3, respectively, and only those fully supported were retained. The second set of genes was parsed in the same way, except for those having one or more exons that failed to align during the first BLASTN analysis. Among these, genes with *N* in their sequence were removed. For the remaining genes, each corresponding genomic locus in the 3 Swe genomes was manually inspected in combination with alignments of short sequencing reads as above, and long reads (ERR3997483, ERR3997394) for Swe1 and Swe3, respectively and only those exhibiting fully supported non-synonymous changes were retained.

## Results

### *D. coniospora* has become more virulent following laboratory culture

We have cultured *D. coniospora* in the laboratory for two decades, passaging it serially through *C. elegans* hundreds of times. We noticed that compared to the original isolate, which we call Swe1, the passaged strain, Swe3, appeared to kill its nematode host more rapidly. As *D. coniospora* can be cryopreserved at -80°C, we were able to compare in the same experiment the Swe1 and Swe3 strains. When we thus assayed the survival of *C. elegans* following *D. coniospora* infection, there was indeed a significant difference between the strains (TD_50_ 40.7 vs 70.8 h for Swe3 vs Swe1, p < 0.0001; Fig. 1A). The speed of killing is influenced by the number of fungal conidia (spores) that attach to the nematode cuticle (Zugasti et al., 2016). Non-infectious spores acquire an adhesive bud as they mature, essential for the capacity to attach to the host. There were no significant differences in spore morphology nor in their rate of maturation between Swe1 and Swe3 (Fig. 1B), with figures that were comparable to those reported previously for *D. coniospora* (van den Boogert et al., 1992). Nor was there any significant difference for spore attachment to *C. elegans* between the two strains (Fig. 1C), suggesting that the observed difference in worm survival was related to other changes affecting fungal virulence. This was supported by the fact that at early time points during the infection, Swe3 provoked a significantly greater host innate immune reaction, as reflected by the higher expression of the *nlp-29p::GFP* transgene (Fig. 1D), a well-characterised reporter of the host response to fungal infection (Pujol et al., 2008). Additionally, there was markedly more rapid growth of mycelia from infected worms before and after they died from infection with Swe3 compared to Swe1 (Fig. 1E). The greater virulence of Swe3 therefore appears to be correlated with an increase in fungal growth during the colonization of *C. elegans*.

**Figure 1:**
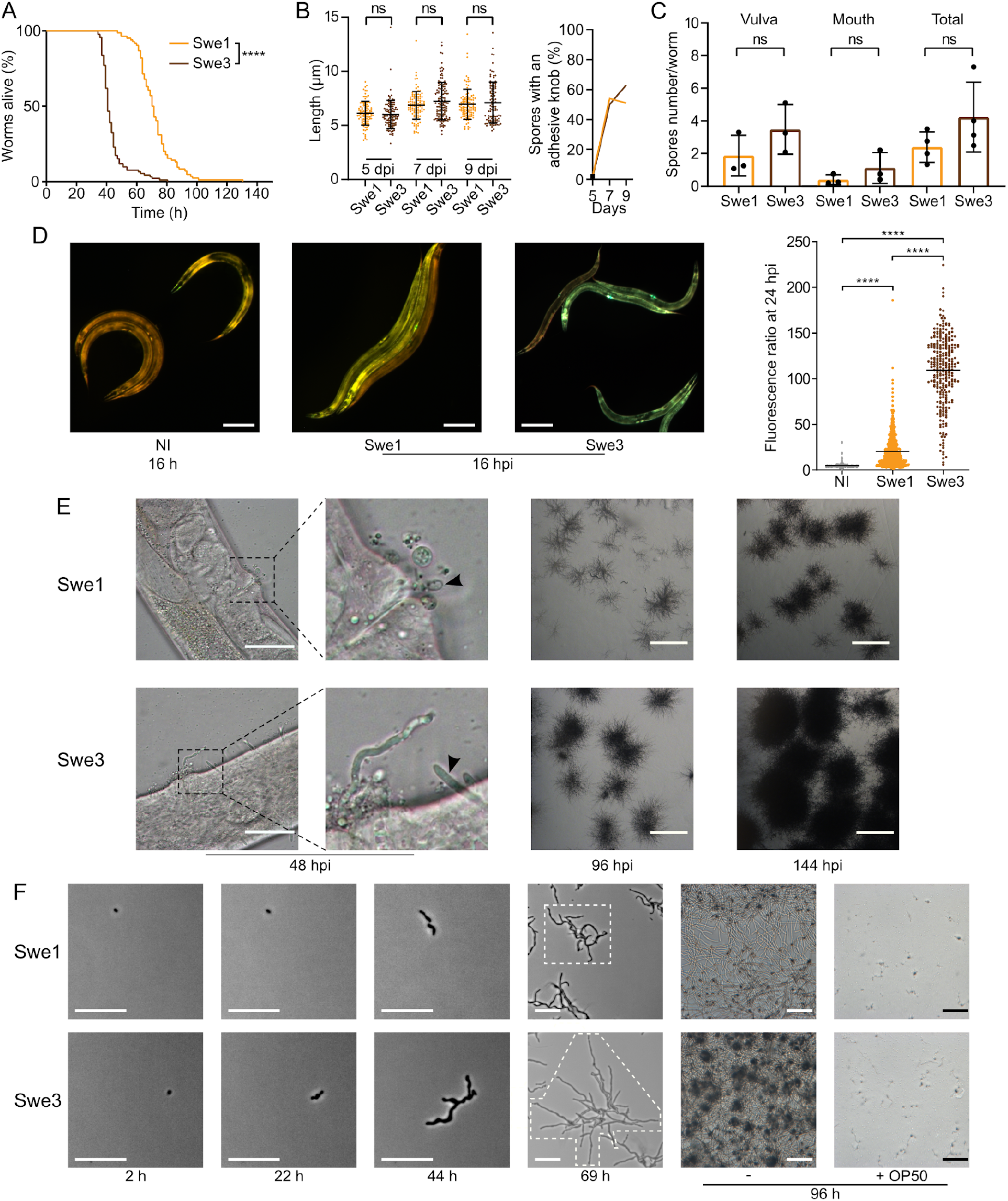
Phenotypic differences between Swe1 and Swe3. (A) Survival of *C. elegans* worms (strain IG463) following infection at 25°C with 1 × 10^9^ spores of Swe1 or Swe3; n = 100 for each strain, **** p < 0.0001, one-sided log rank test. The results are representative of three independent trials. (B) Comparison of the size (left) and prevalence of adhesive buds (right), for spores from Swe1 and Swe3. A minimum of 100 spores were scored at each time point. (C) The number of spores at the mouth and vulval regions of *C. elegans* was counted after 15 h of infection with 1 × 10^9^ spores of Swe1 or Swe3. Each dot represents the mean of an experiment with at least 10 worms; ns = not significant, two tailed unpaired t-test. (D) Left hand panels: Representative fluorescence images of age-matched IG463 worms uninfected (NI) or infected for 16 h (16 hpi) with Swe1 or Swe3. The worm strain carries the integrated transgene *frIs7* that includes *nlp-29p::GFP* and *col-12p::dsRed* transgenes; red and green fluorescence is visualized simultaneously; scale bar: 200 μm. Quantification of the green/red fluorescence ratio (in arbitrary units) of IG463 worms at 24 hpi, compared to aged-matched non-infected (NI) worms; n > 250 worms for each, **** p < 0.0001, unpaired t-test. (E) Representative images of Swe1 or Swe3 infected worms at 48, 96 and 144 hpi. The 2^nd^ panels from the left are a higher magnification of the indicated vulval regions. The arrowhead highlights Swe1 spores attached to the vulva (top), and hyphae growing out from the Swe3-infected worm (bottom); scale bar: 50 μm. The righthand panels show hyphae growing from dead worms; scale bar: 1 mm. (F) Spores of Swe1 or Swe3 (harvested 6 dpi) were seeded on NGM plates, incubated at 25°C without worms and images taken at the indicated times. The extent of hyphal growth from a single spore is delimited by dotted lines in the images at 69 h. The two right-hand columns illustrate the degree of spore germination with or without OP50 after 96 h; scale bar: 100 µm.

To investigate whether the difference in virulence might reflect a more general alteration of fitness, we compared the growth of Swe1 and Swe3 in the absence of worms. These tests were conducted in the absence of bacteria, as we observed that the *E. coli* strain OP50 used with *C. elegans* strongly inhibited fungal growth. Interestingly, Swe3 grew in culture more rapidly than Swe1 (Fig. 1F), suggesting that part of the observed increase in virulence might reflect generally improved growth under laboratory culture conditions, rather than changes affecting specific virulence mechanisms.

### Improving the Swe2 genome sequence

We wished to identify the changes between the Swe1 and Swe3 genomes that could account for their phenotypic differences. The faster growth of Swe3 could reflect changes in mitochondrial function. While the nuclear genome of Swe2, a strain intermediate between Swe1 and Swe3 was sequenced and annotated a few years ago, its mitochondrial genome had not been assembled (Lebrigand et al., 2016). Taking the mitochondrial genomes of Swe1 and Swe3 as a template (Courtine et al., 2020), we remapped the available Swe2 DNA reads, allowing assembly of a complete Swe2 mitochondrial genome. When we compared the genomes of the 3 strains, we found that they were 100% identical. Thus, the phenotypic differences observed between Swe1 and Swe3 must be a consequence of changes to the nuclear genome.

While the Swe1 and Swe3 genomes are more complete than that of Swe2 (Courtine et al., 2020), gene predictions have only been made for Swe2 (Lebrigand et al., 2016). We therefore needed to transpose the Swe2 gene annotations to the other 2 strains’ genomic sequences in order to then be able to compare gene sequences between the different strains. Before doing so, we decided to improve the existing Swe2 sequence and gene predictions. Using the most recent chromosome-level assembly (Courtine et al., 2020) as a starting point, we took advantage of the global synteny between the Swe1 and Swe3 genomes to improve further the assembly of the Swe2 nuclear genome, adding 22 previously un-scaffolded small Swe2 scaffolds, for a total of exactly 580 kb to the existing 31.14 Mb genome (an increase of 0.19%; Supp. Table S3).

At the nucleotide level, the Swe1 and Swe3 genomes are extremely similar, with numerous stretches of >100 kb with 100% sequence identity, the longest being 260 kb. If a particular sequence is absolutely conserved between Swe1 and Swe3, it is reasonable to assume that the Swe2 sequence will be the same. We validated this assumption by amplifying and sequencing 2 randomly selected Swe2 genomic regions containing undetermined (*i*.*e*. “*N*”) nucleotides (Supp. Fig. S2). On this basis, we therefore replaced 19,478 previously undetermined nucleotides in the Swe2 genome with the corresponding sequence from the Swe1/Swe3 genomes, in 471 stretches ranging from 1 to 694 bp (median 23 bp), thereby adding a further 42 kb of defined sequence, removing 7.5 kb of extraneous (erroneously duplicated) sequence, and resulting in an improved sequence for 4 protein coding genes (see Methods for details). This then gave us high-quality chromosomal level assemblies for Swe1, Swe2 and Swe3, that provided a starting point for more detailed analyses.

### Sequence divergence following years of laboratory culture

Unlike the Swe2 genome (Lebrigand et al., 2016), the Swe1 and Swe3 genomes were assembled using long-reads, with short-read polishing. This has the potential to introduce systematic errors, particularly in homopolymeric sequences (Courtine et al., 2020), complicating precise comparisons. Nevertheless, to have a global overview of the sequence divergence between the 3 genomes, we calculated the Average Nucleotide Identity (ANI), a widely applied measure to compare genome sequences, using one alignment-free and two alignment-based methods (Olm et al., 2017). As expected, the sequences were all very similar by this metric (>99.9%). Surprisingly, however, Swe1 and Swe3 showed a higher sequence similarity to each other than to the intermediary strain Swe2 (Supp. Table S4). Therefore, to complement this analysis and provide a more refined view of the sequence divergence between the 3 Swe strains, we aligned their genomes and evaluated the similarity between each pair. We restricted this test to blocks of sequence that gave unambiguous alignments longer than 1 Mbp, to limit the confounding effects of repeated sequences (see Discussion). To obtain an estimate of single nucleotide substitutions, we chose the gap-excluded identity (Li, 2018b). We also used the gap-compressed identity within Minimap2, in which indels of any size are counted as one event. Despite polishing (see above), the Swe2 genome includes stretches of undetermined sequences. We therefore calculated the different metrics with *N* masking when appropriate. The Swe1 and Swe3 genomes were calculated to differ by less than 0.002%, i.e. at one position per 50 kb (Table 1). This value rose to ca. 0.009% when indel events were taken into account. Unexpectedly, but consistent with the results obtained with ANI, the metrics indicated that the genomes of Swe1 and Swe3 were indeed more similar to each other than either were to the genome of Swe2. Compared to the Swe2 genome, the Swe1 and Swe3 genomes differed by 0.004 % and 0.005% respectively. When indels were taken into account, these figures increased, up to 0.06% (Table 1).

**Table 1:** Differences between the Swe genomes

| Query | Subject | Frequency of differences (per Mb) |  |  |  |
| --- | --- | --- | --- | --- | --- |
|  |  | With homopolymers |  | Without homopolymers |  |
|  |  | gap-excluded<br>(Ns removed) | gap-compressed | gap-excluded<br>(Ns removed) | gap-compressed |
| Swe1 | Swe2 | 34 | 550 | 32 | 425 |
| Swe2 | Swe1 | 40 | 512 | 39 | 387 |
| Swe1 | Swe3 | 15 | 83 | 6 | 1 |
| Swe3 | Swe1 | 11 | 86 | 5 | 1 |
| Swe2 | Swe3 | 45 | 512 | 30 | 400 |
| Swe3 | Swe2 | 47 | 550 | 32 | 412 |

Inspection revealed that many instances of potential differences occurred in stretches of homopolymeric sequence. It is known that correctly calling homopolymers from long-reads is challenging. Our analysis indicates that even with short-read polishing (Courtine et al., 2020), the Swe1 and Swe3 genomes contain residual homopolymer errors (Supp. Fig. S3). To have a more realistic measure of sequence similarity, we therefore removed homopolymer sequences of 5 or more identical nucleotides, from each of the Swe genomes, and recalculated the different metrics. In all comparisons, the similarity scores increased, with a more marked effect for comparisons involving Swe2 (Table 1). Nevertheless, Swe1 and Swe3 were still reported to have the highest similarity, with a divergence as low as 0.0005% and 0.0001% (i.e. one change per 200 kb and 1 Mb) by the gap-excluded and gap-compressed metrics, respectively. The values for the comparison with Swe2 were an order of magnitude higher (0.004% and 0.003% of divergence with Swe1 and Swe3, respectively), and about 0.04% of divergence using the gap-compressed metric. As discussed below, this likely reflects the different genome sequencing approaches. Nevertheless, the values for Swe1 and Swe3 give an estimate of the comparatively low sequence evolution of the Swe genomes over time.

### No sexual reproduction in *D. coniospora* under laboratory conditions

Any interpretation of *D. coniospora* molecular evolution requires a knowledge of its reproductive mode. Whether or not *D. coniospora* has a sexual cycle remains unclear. Sexually reproducing fungi can be homothallic (self-fertile) or heterothallic (outcrossing) (Heitman et al., 2007). In ascomycetes, this is determined by the allelic combination of the genes of the mating type (*MAT*) loci, termed idiomorphs. In a homothallic species, individual cells will possess both *MAT1-1* and *MAT1-2* idiomorphs, while in heterothallic species, a single cell will have genes of only one idiomorph. Mating is then restricted to cells with complementary idiomorphs. Zhang *et al*. reported the *D. coniospora* ARSEF 6962 (called here Dan2) genome to include a well-conserved *MAT1-1* locus composed of 3 genes (*MAT1-1-1* (KYK59754.1), *MAT1-1-2* (KYK59755.1) and *MAT1-1-3* (KYK59756.1)), but no *MAT1-2* idiomorphs. Taken together with evidence for an active repeat induced point mutation (RIP) system, and the presence of genes for 2 RIP system proteins, as well as 4 heterokaryon incompatibility (HET) proteins, they suggested that *D. coniospor*a might be heterothallic (Zhang et al., 2016). Dan2 is a derivative of the *D. coniospora* isolate CBS 615.82 that we refer to as Dan1 (Courtine et al., 2020). Like Dan2, the Dan1 genome contains genes for the 2 RIP system and 4 HET proteins, as well as 3 *MAT1-1* genes in a single locus, but no *MAT1-2* genes. While the Swe genomes contain genes for the same RIP system and HET proteins (Supp. Table 5), no *MAT* genes were predicted in the original annotation of the Swe2 genome (Lebrigand et al., 2016). On the other hand, in the Swe2 region syntenic to the Dan1/Dan2 *MAT1-1* locus, using BLASTX and TBLASTN searches, we found a putative *MAT1-2-1* gene that had not been previously annotated, fully conserved in Swe1 and Swe3, but absent from the Dan1 and Dan2 genomes (Fig. 2). Zhang *et al*. had proposed that *D. coniospora*’s adaptation to an endoparasitic lifestyle might result in a gradual loss of sexual reproduction (Zhang et al., 2016). On the basis of our analysis, while it remains possible that in nature *D. coniospora* is heterothallic, under our experimental conditions, the Swe strains used here will only reproduce in an asexual manner. Since spore formation involves a transition between two haploid states, with no meiosis, for this study, the *D. coniospora* isolate Swe1 and its derivatives can be considered to behave as haploid asexual clones.

**Figure 2:**
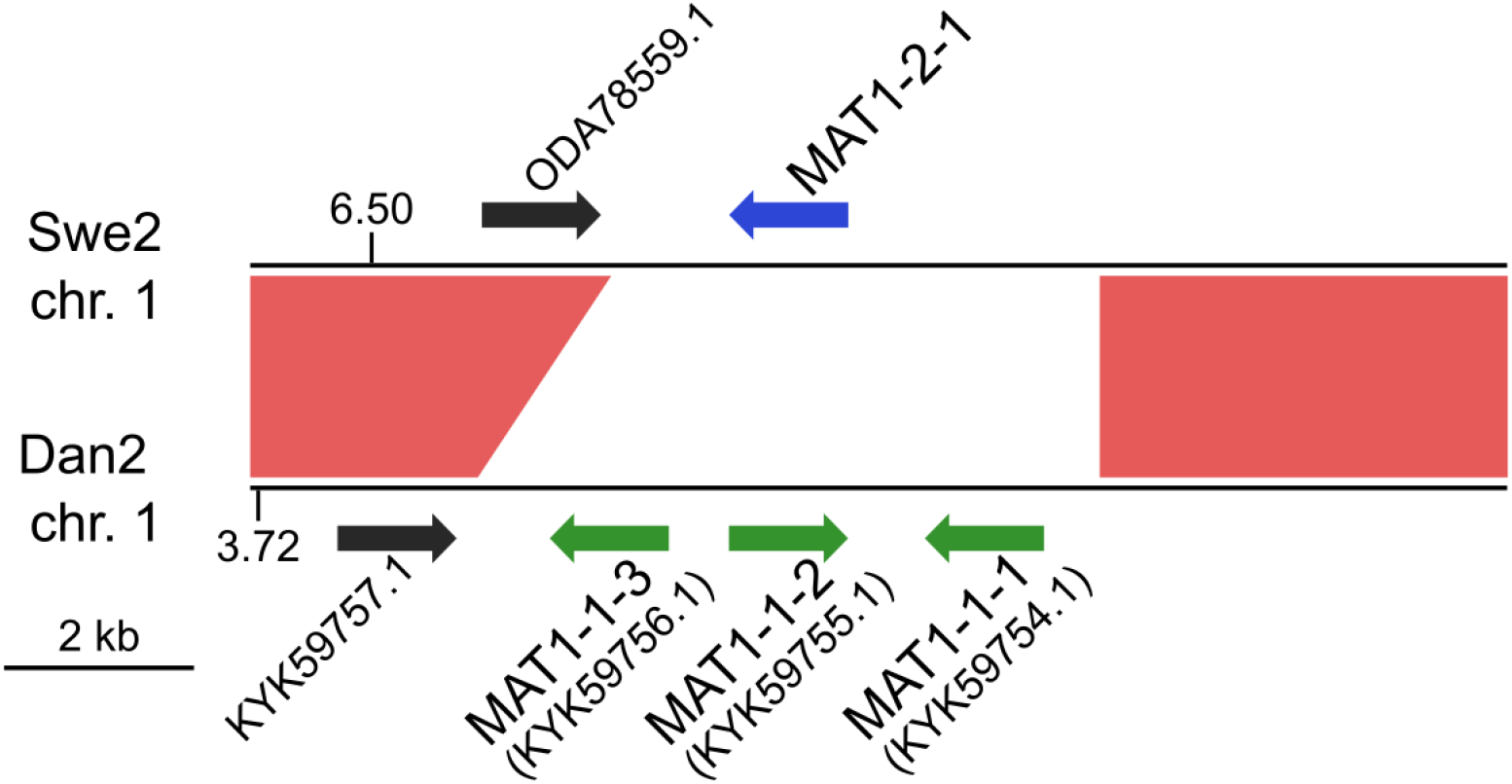
MAT loci in *D. coniospora*. The Swe2 region syntenic to the Dan2 MAT locus is shown. The previously annotated Dan2 MAT genes are indicated with green arrows, labeled with their idiomorph identifiers. The newly identified Swe2 *MAT1-2-1* gene is shown in blue. The red shapes highlight the extent of the sequence synteny, and the black arrows represent a pair of neighbouring orthologous genes.

### Completion of the Swe proteome

The absence of *MAT1-2-1* from the predicted gene set for Swe2 suggested that there might be other lacunae in gene prediction. We therefore conducted an all-against-all comparison between the protein-coding genes of Dan2 and Swe2 (Genbank GCA_001625195.1 and GCA_001618945.1, respectively). This identified 298 genes present in Dan2 for which there was no equivalent prediction for Swe2. Detailed sequence searches in the Swe2 genome (see Methods) revealed plausible sequences for 39 of them, and these were added to the Swe2 gene set. This set was then used to predict genes in the Swe1 and Swe3 genomes, using a sequential approach (see Methods). Direct searches of the remaining 259 Dan2 genes against the Swe1 genome failed to identify any further previously unpredicted genes.

### Genomic comparison of the 3 *D. coniospora* strains reveals mutated genes

We then compared the respective sequences of the complete set of 8,702 protein-coding genes from the Swe1, Swe2, and Swe3 genomes. The vast majority, 94.7%, had precisely the same DNA sequence in the 3 genomes (Fig. 3). The remaining 457 genes potentially contain mutations. In most cases (68.4%), the mutation was present in an intron. In no case did this affect splice donor or acceptor sites and these mutations would not be expected to affect the corresponding protein. We filtered out 103 of the remaining 144 genes either because the mutation was synonymous, and would not result in any change in the sequence of the corresponding protein, or since a potential mutation had been called because the polished Swe2 sequence still contained one or more *N*. Even though the 3 genomes are of high quality, 31 out of the remaining 41 potential mutations were found to be a consequence of an error in a homopolymeric sequence, despite short-read support for the consensus sequence (e.g. Supp. Fig. S3). These errors in the Swe1 and/or Swe3 genomes were manually corrected. It should be noted that similar errors will exist in non-coding regions that were not examined here. In the end, we identified with high confidence just 10 protein-coding genes with indels or non-synonymous mutations in exonic sequences.

**Figure 3:**
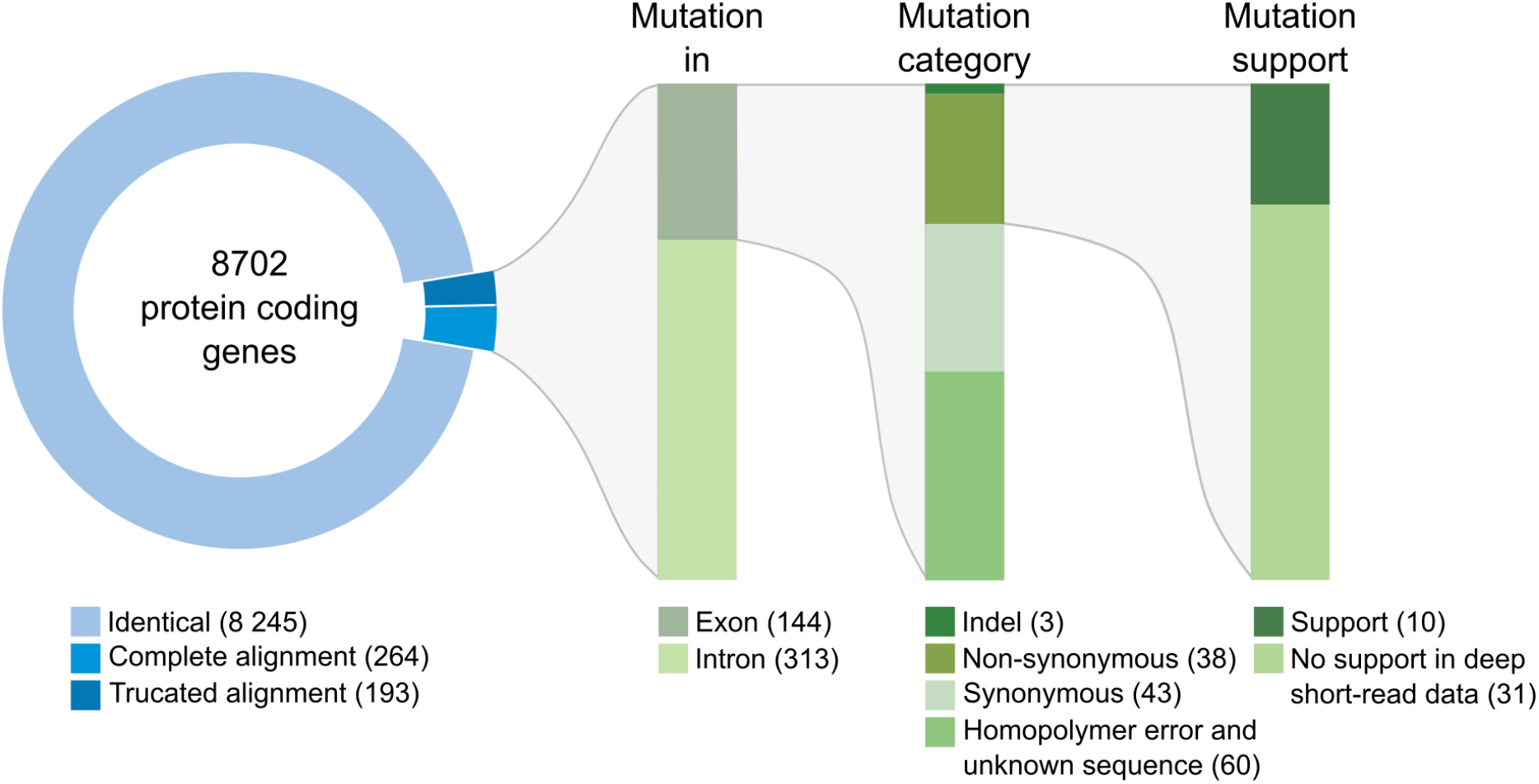
Classification of mutations identified between the genomes of the Swe strains. The overview includes the number of genes and mutations identified at each step of the workflow (see Methods for details). Mutations identified in introns were individually examined; none affected splice donor or acceptor sites.

When applicable, the changes that we found in the genomic sequence were fully supported by the transcriptomics data available for Swe2. Interestingly, in two cases, specifically for mutations that were predicted to appear between Swe2 and Swe3, the RNAseq reads were indicative of sequence heterogeneity (Supp. Fig. S4). The samples for RNAseq analysis had been collected following a particular culture regime, involving amplification in liquid media (Lebrigand et al., 2016). It appears that the RNAseq captured a snapshot of a heterogeneous population, with a mixture of both mutated and non-mutated fungal lineages, prior to subsequent fixation of the respective mutations.

The first example where we captured a mutation before it became the preponderant allele in the population corresponds to g932.t1/ODA82425.1 that encodes a transcription factor related to sterol regulatory element binding proteins (SREBP), homologous to *Sre1* in the fission yeast *Schizosaccharomyces pombe*. Swe1 and Swe2 share a common sequence, while in Swe3, the gene carries a premature stop codon that would be predicted to be a null mutation (Table 2). *Sre1* is activated under hypoxic and sterol-depleted conditions and upregulates the expression of genes involved in sterol biosynthesis (Hughes et al., 2005). A similar process exists in *Aspergillus* species, for which the ability to grow under hypoxic conditions is linked to virulence (Chung et al., 2014). In fungi of the classes *Sordariomycetes* and *Leotiomycetes*, relatively close phylogenetically to *Drechmeria*, the homologues of SREBP, such as *FgSre1* in *Fusarium graminearum*, are required for hypoxic growth, but are not involved in the regulation of sterol biosynthesis, which is controlled by the sterol uptake control protein *FgSR* (Liu et al., 2019). The gene g932.t1, which we refer to here as *DcSre1*, is therefore potentially involved in growth under hypoxic and/or sterol-depleted conditions.

**Table 2:**
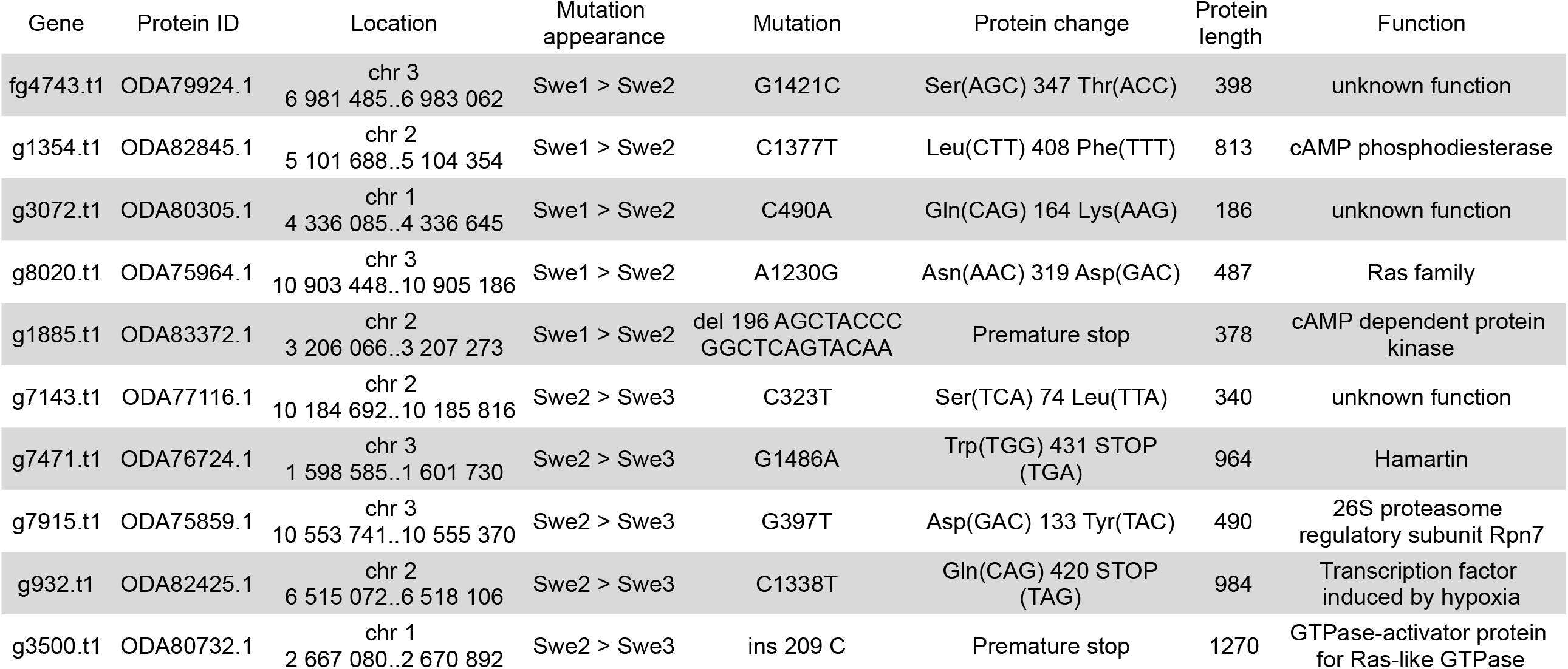
Summary of non-synonymous mutations

While in natural environments sterols can be limited, to support *C. elegans* growth, NGM is supplemented with cholesterol. The loss of *DcSre1* might have arisen as a consequence of prolonged growth on a sterol replete medium. To address the potential function of *DcSre1*, we therefore first assayed the growth of Swe3 on agar plates of NGM with or without added cholesterol. We observed no obvious difference between the high and low cholesterol conditions (Supp. Fig. S5). Since even without *DcSre1*, Swe3 is able to maintain its growth regardless of sterol levels, this suggests that *DcSre1*, like *FgSre1*, is not involved in the regulation of sterol biosynthesis. *D. coniospora* does possess an orthologue of *FgSR* (ODA81938.1), and we hypothesise that this plays the role of sterol biosynthesis regulator as in closely-related species. On the other hand, when we assayed Swe1 and Swe3 in the presence of CoCl_2_, which mimics hypoxia (Lee et al., 2007), there was a significant difference in the growth of the two strains. While under standard conditions Swe3 grew faster than Swe1 (Fig. 1F, Fig. 4A), at a concentration of 1 mM of CoCl_2_, the situation was reversed and Swe1 clearly grew better than Swe3. Indeed Swe3’s growth was as strongly inhibited as it was in the presence of 2 mM CoCl_2_, when Swe1 failed to grow too. Thus, presumably as a consequence of the mutation in *DcSre1*, Swe3 has lost the capacity to adapt to conditions that mimic hypoxia. Whether loss of *DcSre1* function also leads to better growth under laboratory conditions, both in the absence of worms and during infection, remains to be determined.

**Figure 4:**
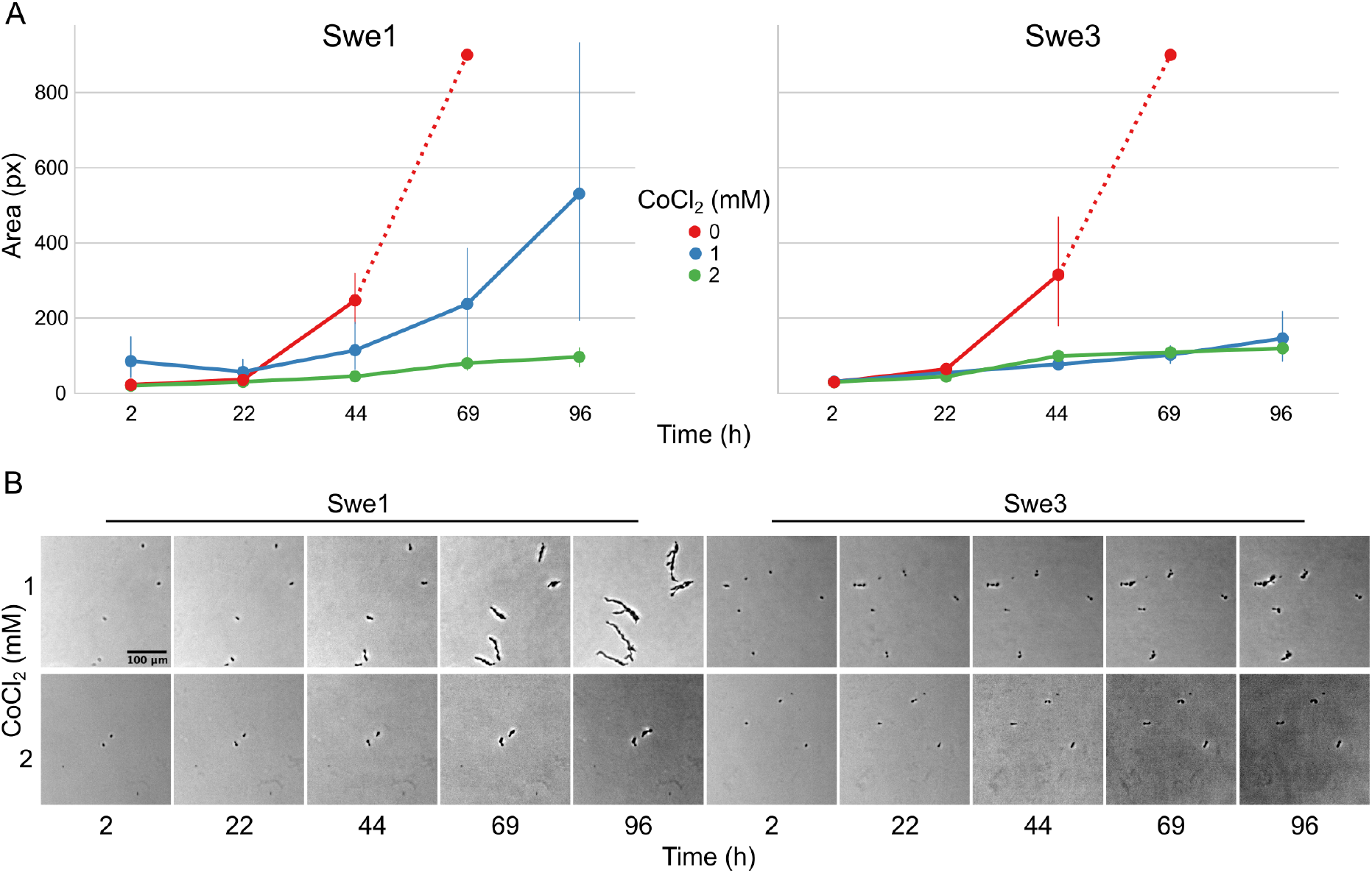
Comparison of the growth of Swe1 and Swe3 in hypoxia-like conditions. (A) Quantification of growth rates for Swe1 and Swe3 on standard NGM plates (0 mM), or plates supplemented with CoCl_2_ to a final concentration of 1 or 2 mM. As illustrated in Figure 1F, measurement beyond 44 h of Swe1 and Swe3 growth in the absence of CoCl_2_ is confounded by the overgrowth of mycelia derived from separate spores. The average size and 95% confidence interval for 8 - 35 individual spores per condition, followed at each time point, are shown. (B) Representative images of spores of Swe1 and Swe3 growing in the presence of 1 or 2 mM CoCl_2_. A direct comparison can be made with Figure 1F, as the experiments were conducted in parallel.

The second gene for which there was evidence for the mutation arising during liquid culture of Swe2 and being fixed in Swe3 was g7915.t1/ODA75859.1 (Table 2). It corresponds to the RPN7 subunit of the proteasome lid, required for the structural integrity of the complex (Isono et al., 2004). We identified a non-synonymous mutation at a highly conserved position within the Pfam *RPN7* (PF10602) domain. The proteasome serves an essential role in cell physiology. We would predict this mutation to affect overall organismal fitness rather than being directly important for virulence.

Several other mutations affect genes involved in important cell functions (Table 2). Thus, we identified a nonsense mutation that occurred between Swe2 and Swe3 in the gene g7471.t1/ODA76724.1, encoding a protein similar to hamartin/tuberous sclerosis protein 1 (TSC1). The TSC1/TSC2 complex has GTPase Activating Protein (GAP) activity and is known to inactivate the Ras GTPase, Rheb (Rhb1/a). In *S. pombe*, loss of Tsc1 and/or Tsc2 function results in a decreased uptake of arginine and impacts amino acid biosynthesis (van Slegtenhorst et al., 2004). A second mutation, in gene g8020.t1/ODA75964.1, also potentially impacts Ras signalling as a non-synonymous change, N319D, occurred between Swe1 and Swe2 in the *D. coniospora Ras* gene itself. The mutation is predicted to affect the interaction with the Ras regulator GDP Dissociation Inhibitor that stabilises the inactive (GDP-bound) form of Ras in the cytosol (Müller and Goody, 2018). The mutations observed in *DcTsc1* and *DcRas1* have the potential to affect the overall growth capacity of *D. coniospora*.

Similarly, the predicted null mutation in gene g3500.t1/ODA80732.1 (a single nucleotide insertion causing a frameshift and so a precocious stop codon) affects the *D. coniospora* Bud2p homologue. In *S. cerevisiae*, Ras-GAP Bud2p activates the Ras protein Rsr1p/Bud1p, which controls the site of budding. Loss of function of *Bud2* leads to the constitutive activation of Rsr1p/Bud1p and thus to a random budding site (Ni and Snyder, 2001). In *D. coniospora*, loss of DcBud2 might be expected to affect the pattern of hyphal branching, but the ramifications from a single germinating spore (see for example Figs. 1E, 4B) confounded attempts at quantitative comparison. Nevertheless, again, this is a mutation more likely to affect growth than virulence specifically.

The remaining mutations affecting genes with well-characterised orthologues correspond to a pair of proteins that potentially function in the same regulatory pathway (Table 2). For one, gene g1354.t1/ODA82845.1, a non-synonymous mutation, L408F, occurred between Swe1 and Swe2. This gene (*DcPde1*) is predicted to encode a 3’ 5’-cAMP phosphodiesterase (PDE). Loss of PDE function would be expected to result in increased cAMP levels. For the second, gene g1885.t1/ODA83372.1, a 20 bp duplication in the Swe1 genome, predicted to render the corresponding protein non-functional, was lost in Swe2. Thus in this case, the mutation would be expected to restore function to the predicted regulatory subunit of a cAMP-dependent protein kinase (*DcCPKA*). In many fungi, cAMP levels act via cAMP-dependent protein kinases to influence growth, morphology and sporulation (Kim et al., 2011; Wang et al., 2011). Assuming that the mutation of *DcPde1* is a loss-of-function, in combination with the mutation in *DcCPKA*, growth and sporulation in Swe2 and Swe3 would be predicted to be more tightly controlled by the ATP/cAMP balance (Fig. 5).

**Figure 5:**
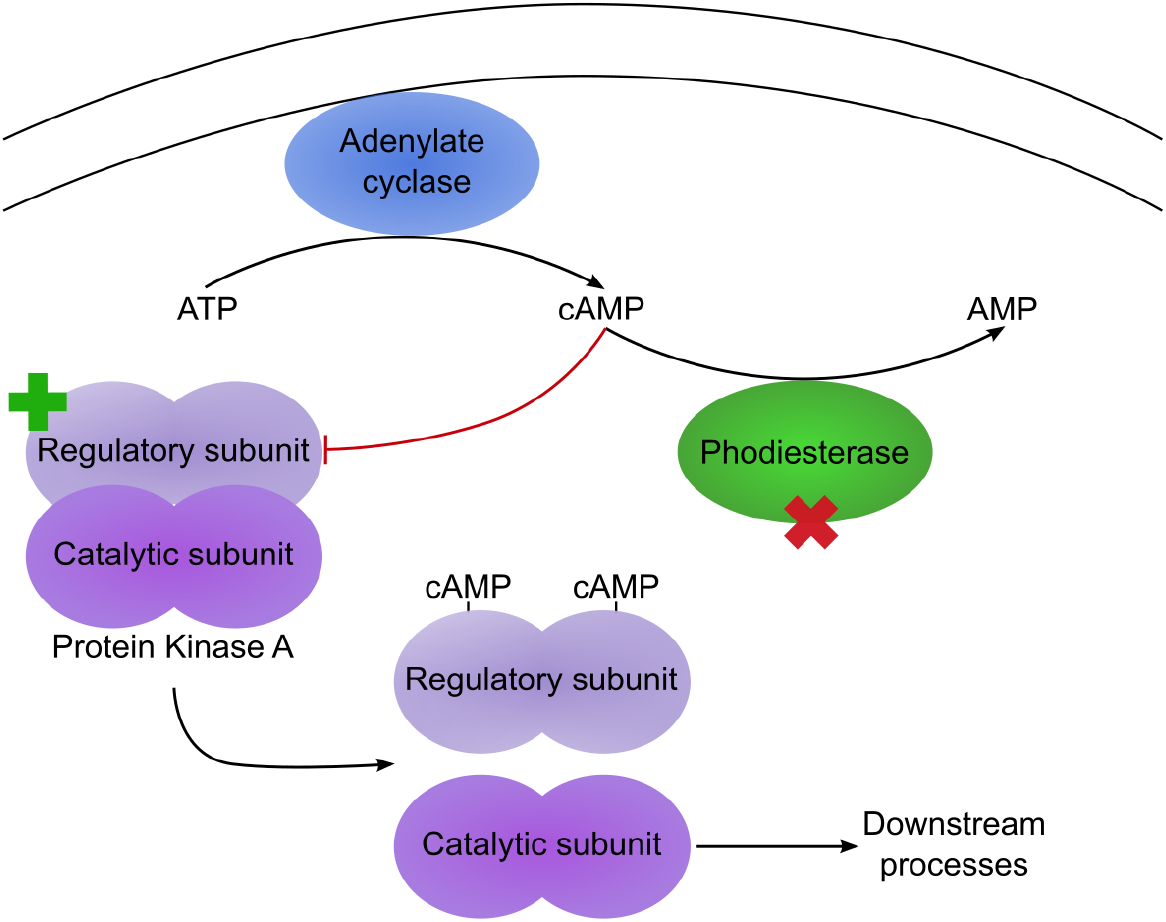
Two mutations potentially affect the same pathway. Simplified diagram illustrating the regulation of protein kinase A (PKA) by cAMP. Under low cAMP condition, the PKA regulatory and catalytic subunits dimerize, leading to an inactive PKA. In Swe1, as the regulatory subunit is constitutively inactive, PKA would be predicted to be permanently active. In Swe2, the inactivating mutation of the regulatory subunit is reverted to wild-type (as indicated by the green cross), so PKA would again come under the control of cAMP levels. Its activity is also regulated by a cAMP phosphodiesterase (PDE), as this converts cAMP to AMP. The PDE bears a non-synonymous mutation in Swe2 (red cross). Assuming that this corresponds to a loss of function, the two mutations would act synergistically, as cAMP would accumulate, thereby accentuating the suppression of repression of PKA activity. Thus PKA would be predicted to be more active in Swe2 (and Swe3) than in Swe1 under the same culture conditions.

The remaining 3 proteins have poor or non-existent annotation. Genes g7143.t1/ODA77116.1 and g3072.t1/ODA80305.1 that acquired non-synonymous mutations at different steps of culture (Table 2) correspond to hypothetical proteins of unknown function, found in a phylogenetically restricted range of fungi. Still more extreme is the case of fg4743.t1/ODA79924.1. Despite using sensitive search tools (including HHblits (Steinegger et al., 2019) against the UniRef100 database), we were unable to find any plausible homologues in any other species. Indeed, there is no equivalent of fg4743.t1 even in the genomes of the 2 available *D. coniospora* Dan strains. The gene was predicted *ab initio* with the software tool Augustus (Lebrigand et al., 2016), but has only very marginal RNAseq read support, with 11 and 16 reads in samples from mycelia and spores, respectively, not covering all of the predicted coding sequence and with none of the reads confirming the predicted intron-exon boundaries (Supp. Fig. S6). We were unable to amplify a transcript from cDNA using sequence-specific PCR primers designed on the basis of the existing gene model. Thus if such a gene does exist, it is unique to the Swe strains of *D. coniospora* and its sequence gives no clue as to its possible function. Like all the genes described above, its potential role in virulence *per se* remains to be addressed.

## Discussion

The difference in virulence between Swe1 and its domesticated derivative Swe3 opened a potential path to dissect the molecular basis of pathogenicity in *D. coniospora*. As mentioned above, at the outset, we had little idea that we would culture *D. coniospora* for 20 years. Otherwise, we would have made adjustments to our method. We used the wild-type N2 strain of *C. elegans* to propagate *D. coniospora*. In line with recommended practice, the cultures of N2 were replaced from frozen stocks several times a year. Although this was not the specific intention, as a consequence, *D. coniospora* was grown in a host with an effectively stable genetic background. While this simplifies interpretation of any changes to *D. coniospora*, had a replicate been included for which N2 was not replaced, but allowed to accumulate mutations, we might have gained insight into host-pathogen co-evolution.

Our *in silico* analysis was, therefore, necessarily restricted to a genomic comparison of the different *D. coniospora* strains. As a first step, we improved the genome assembly for Swe2, as this served as our reference sequence for gene curation in the other two strains. The refined Swe2 genome is of high quality, with less than 1/1000 undefined bases on average. Given the degree of divergence in long fully-defined regions of the Swe genomes (i.e. 0.5 substitutions per 100 kb between Swe1 and Swe3), one would expect < 70 changes in exonic sequence. Thus, we can predict that few if any SNPs and indels affecting protein-coding genes have been overlooked because of the remaining undefined bases. In the context of this study, improving the genome sequences further would require a disproportionate effort for relatively limited potential return.

Unexpectedly, on the basis of standard similarity metrics, the Swe1 and Swe3 sequences were reported to be more similar to each other than to Swe2, which makes little biological sense. This was also the case when the effect of erroneous homopolymer length determination associated with nanopore sequencing was eliminated. This difference is likely to reflect the strategy used to assemble the Swe1 and Swe3 genomes, compared to that for Swe2. Firstly, the average short-read coverage for Swe1 and Swe3 was close to 400X, four times higher than that for Swe2. This should increase the intrinsic sequence accuracy at each position. Perhaps more importantly, the long reads used for assembling the Swe1 and Swe3 genomes allowed unambiguous positions to be assigned to repeated elements, including transposons. During polishing, short-reads were recruited to the correct sequence with high-fidelity. For the Swe2 genome, without the support of long reads, some repeated elements were mis-positioned, or had an incorrect copy number, leading to genome compression, as described previously (e.g. (Eccles et al., 2018)), and reflected in the slightly smaller (ca. 1%) total size for the current Swe2 genome. In the absence of correct assembly, some short reads would be mapped to an erroneous genomic region and this recruitment of reads from one or more slightly divergent copies of the same element would lead to inaccuracies in the final sequence. We indeed observed this type of event in our manual inspection of short-read mapping to the Swe2 genome. This is likely to be a common artifact when comparing genomes sequenced using different technologies.

The mutations that accumulated in Swe3, compared to Swe1, were far from randomly distributed. Introns and exons represent about 5% and 40% of the genome sequence, respectively. Given a divergence rate of about 0.002% between Swe1 and Swe3, one would expect around 8 and 60 nucleotide changes in introns and exons, respectively. This is very far from our estimates (incorporating the measured true positive rate) of 78 and 20, respectively, presumably a reflection of selection pressure maintaining the integrity of protein-coding genes.

At the DNA level, we found no evidence for incomplete spread of the different alleles that accumulated in Swe2 and then Swe3. In the simplest case of haploid selection, as a very rough estimate, a new allele that confers a 10% fitness advantage can go from a low to a high frequency in a population in 100 generations, while one associated with a 1% increase in fitness will take 1000 generations. For this experiment with *D. coniospora*, a generation can be equated to one passage, from the addition of spores to a plate of *C. elegans*, to the harvesting of new spores, typically 1-2 weeks later. We have no way of knowing exactly when each new allele arose, but given their spread in a relatively small number of generations (estimated to be 25/year on average), excepting any case of hitch-hiking by a neutral allele, each is likely to confer a substantial increase in fitness. Ideally, we would be able to introduce the alleles singly into the Swe1 background (or that of Swe2, as appropriate) to determine their individual contributions to fitness and/or virulence. Our established *D. coniospora* transformation method (He and Ewbank, 2017) no longer functions and we are not currently able to alter specifically the fungal genome. As an alternative, one can heterologously express candidate virulence factors in *C. elegans* (Zhang et al., 2021), but this is only appropriate for *D. coniospora* proteins that are predicted to be secreted into the host, which is not the case for any of the candidates identified in the current study.

The most striking molecular change concerned *DcCPKA*. In Swe1, the gene is predicted to be non-functional, due to a tandem duplication of a short sequence element. This was lost in Swe2, putatively restoring function. Similar effects have been reported following long-term culture of *S. cerevisiae*, including one linked to adenine biosynthesis (*ade2-1*), wherein mutations reverted a premature stop codon in the parental strain so that the full ADE2 sequence could be translated (Johnson et al., 2021). Rearrangements involving tandem sequence elements like this can arise due to replication slippage and are well-known drivers of genomic change (Hancock, 1996). In the Dan1 and Dan2 genomes, the *DcCPKA* locus resembles that of Swe2 and is thus predicted to encode a functional protein. Sequence analysis of other environmental isolates of *D. coniospora* will be needed to establish how often *DcCPKA* function is lost in nature. It would also be extremely interesting to know whether the *DcCPKA* mutation appeared in our laboratory culture before or after the mutation in *DcPde1*, but unfortunately we have no cryopreserved samples between Swe1 and Swe2. Since we cannot be certain that the *DcPde1* mutation in Swe2 represents a loss of function allele, it is also an open question as to whether the 2 mutations, in *DcCPKA* and *DcPde1*, are compensatory or mutually reinforce an effect on PKA signaling. Our current model is that the Swe2 *DcCPKA* allele encodes a functional regulatory protein, subject to control by cAMP levels, and that with decreased DcPde1p activity, the PKA pathway is more repressed in Swe2 than Swe1 for given levels of ATP and adenylate cyclase activity. This model could be tested by site-specific mutation of the Swe1 genome, but, unfortunately, as explained above, this is currently not possible. Indeed, this is an important barrier to making definitive causative links between any of the observed mutations and the observed alteration of virulence.

Another potential limitation of our study reflects the use of Swe2 gene set as our reference. While we found no more paralogues for Dan2 genes in the Swe1 genome than in the Swe2 genome, if there are genes with mutations substantially altering their predicted coding sequence (e.g. introduction of a premature stop codon) in Swe2, they would be missing from the gene set and therefore not included in our analysis. Nevertheless, in common with *DcCPKA* and *DcPde1*, several of the mutations that were identified between Swe1 and Swe3 are likely to play general roles in fungal growth. Compared to the natural environment, laboratory culture conditions are relatively constant and it may be advantageous for *D. coniospora* to lose regulatory pathways that are not needed, as has been seen in other evolution experiments (reviewed in (Kvitek and Sherlock, 2013; McDonald, 2019). Indeed, in one replicated study, a majority of mutations were found in three major signaling networks that regulate growth control, including the Ras/cAMP/PKA pathway that in *D. coniospora* involves *DcCPKA* and *DcPde1*. Although such mutations are predicted to be beneficial in a stable environment, they generally come at the cost of eliminating the metabolic plasticity needed when nutrient supplies change. We assume that this type of mutation will impact pathogenesis through a general effect on growth. It remains an open question whether any of the other mutations we identified in 3 unannotated predicted hypothetical proteins might impact virulence *per se*. As they are not predicted to be secreted, they are unlikely to be direct effectors of fungal virulence, but could conceivably be involved in the regulation of virulence factor gene expression. In the absence of tools for site-directed mutagenesis, determining their function remains a challenge for the future.

## Data Availability

All relevant data can be accessed either at the EBI/NCBI/DDBJ via BioProject PRJNA269584, or on our institutional server at http://www.ciml.univ-mrs.fr/applications/DC/Genome.htm.

## Acknowledgements

We thank Marie-Anne Félix for insightful comments, Philippe Fort for helpful discussion concerning Ras GAP function and phylogeny, and the CIML imaging (ImagImm) and bioinformatics platforms.

---

**Supplementary Figure S1:**
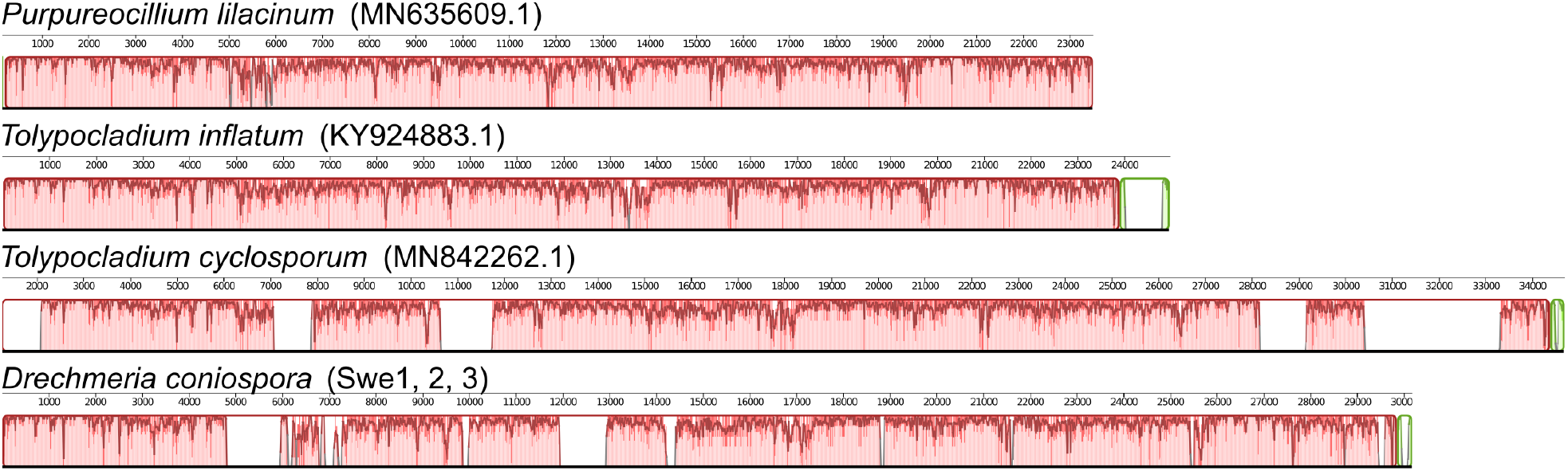
Alignment of mitochondrial genomes from *D. coniospora* and 3 closely related fungi. For each species, a histogram of the overall sequence conservation (from 0 to 100%) calculated using a sliding window of fixed size is shown under the genome coordinates (in kb). The regions in red are syntenic and conserved across the four species, those in green correspond to sequences absent from *P. lilacinum*.

**Supplementary Figure S2:**
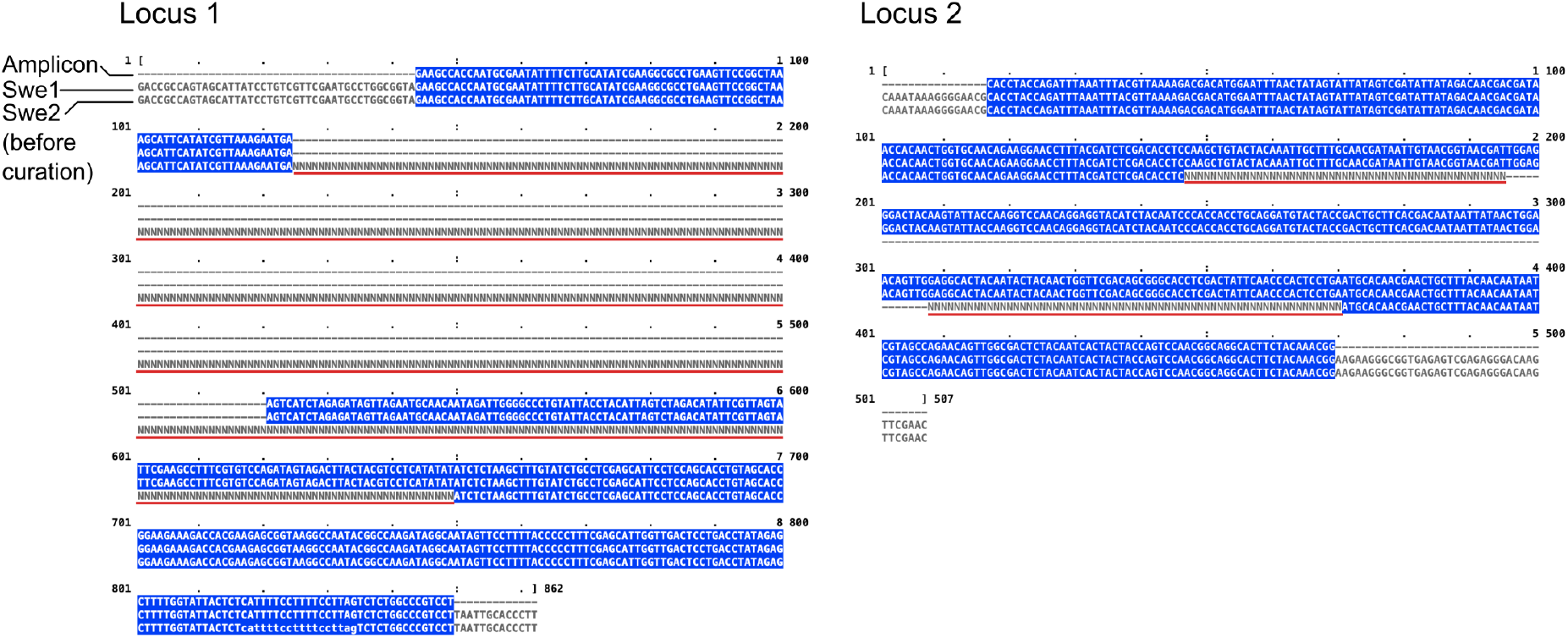
Experimental validation of strategy to replace undetermined Swe2 sequences. Sequence alignments of 2 sequenced amplicons from Swe2, and the corresponding regions from the Swe1 and existing Swe2 genomic sequences. The blue highlights matching positions between the 3 sequences. The red lines indicate the stretches of underdetermined bases in the original Swe2 genome. Note, these regions were chosen as the corresponding sequence in Swe3 is identical to that in Swe1. The primers pairs used were GACCGCCAGTAGCATTATCC and AAGGGTGCAATTAAGGACGG for the left locus; CAAATAAAGGGGAACGCACC and GTTCGAACTTGTCCCTCTCG for the right.

**Supplementary Figure S3:**
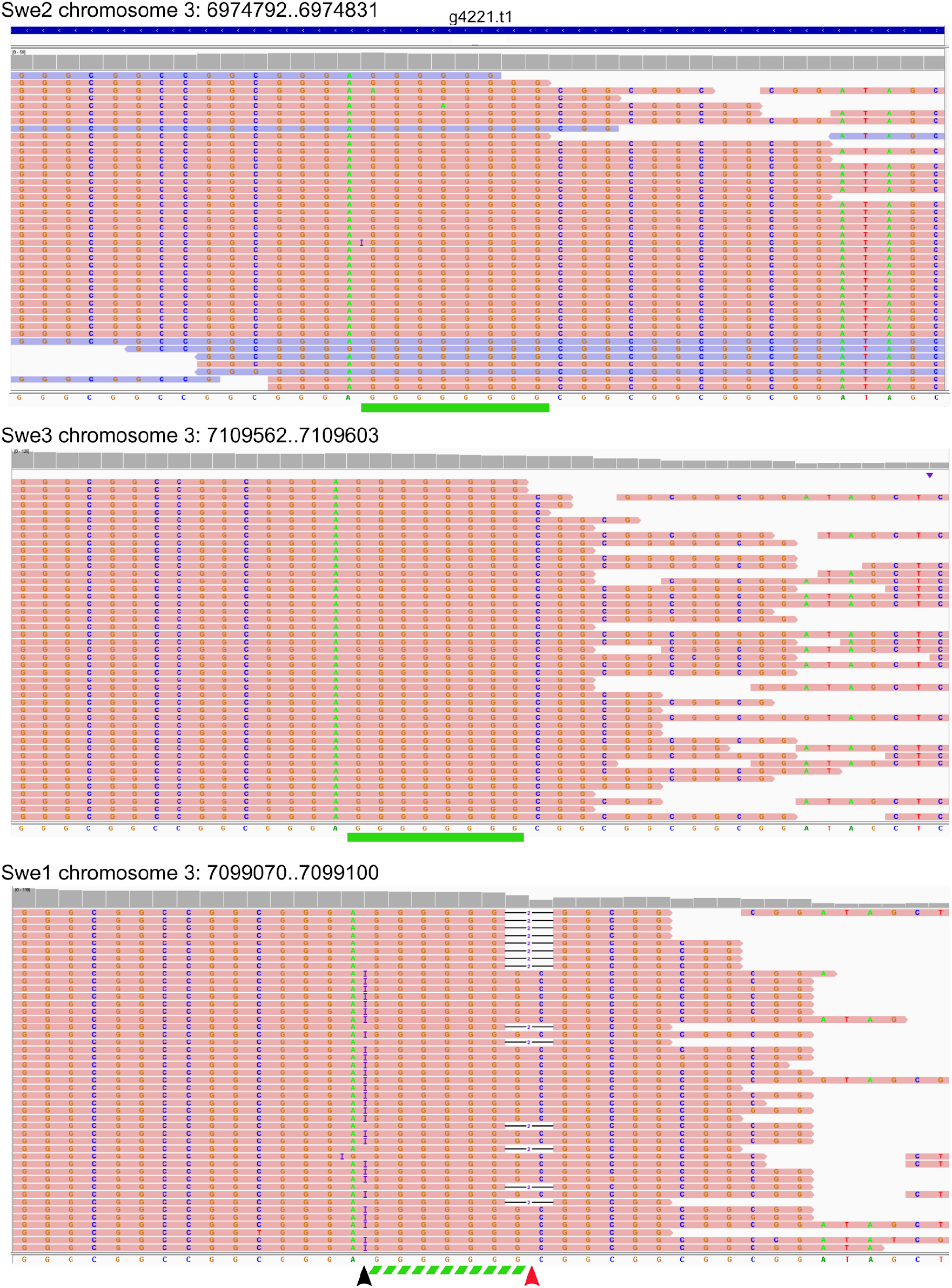
Homopolymeric region bearing a residual sequence error. Mapping of short DNA sequencing reads to the equivalent locus in the Swe2, Swe3 and Swe1 genomes, for which the respective genomic coordinates are indicated. A selection of the reads for each is represented, with their sequences in a salmon or light blue background, depending on their orientation. The sequence predicted for the locus, after polishing for Swe1 and Swe3 (Courtine et al., 2020), is shown below the mapped reads in each panel. The green rectangles under the top and middle panels highlight a homopolymer of 8 Gs. This 8 G sequence, supported by a large majority of short reads in all cases, has been incorrectly truncated to 7 Gs in Swe1, as highlighted by the dashed green rectangle. The black arrowhead marks the position where numerous reads have an extra G (an insertion as indicated by the blue I) that has not been taken into proper consideration, and the red arrowhead the highlights a position where numerous reads have been mismapped because of the tandem repetition of the sequence CGG, 3’ to the homopolymeric sequence, resulting in the inference of 2 nucleotide deletions, with respect to the (erroneous) reference, as indicated by -2-.

**Supplementary Figure S4:**
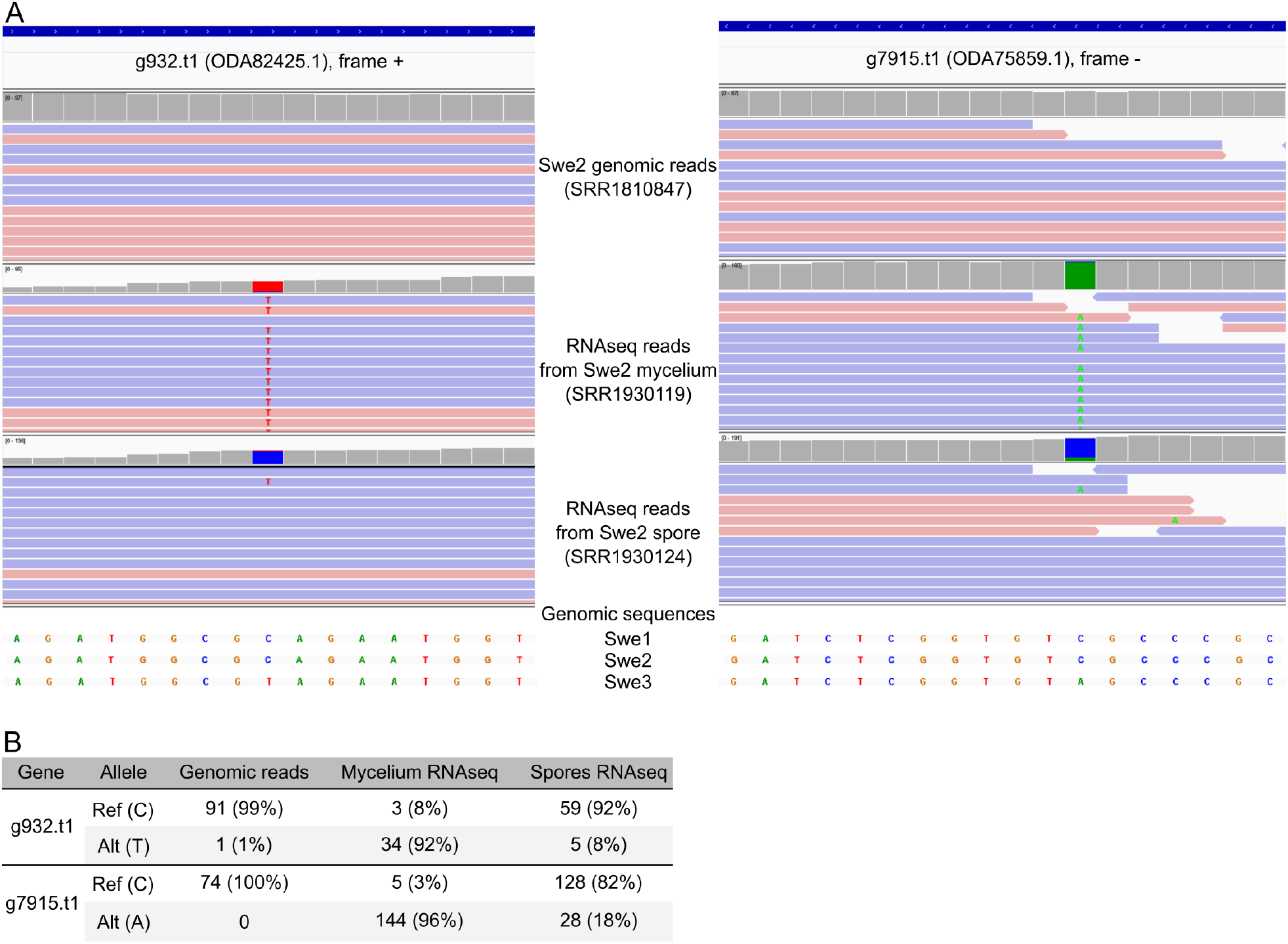
Examples of sequence heterogeneity captured by RNAseq. (A) Evidence at two sites for a disparity between the Swe2 genomic sequence and the sequence determined by transcriptome sequencing. Screen captures from IGV showing the mapping of genomic reads (top), and two RNAseq read sets from mycelia (middle) and spores (bottom). The boxes at the top of each panel represent the read coverage at each nucleotide position. They are coloured grey if >95% of reads support the consensus sequence for the Swe2 genome (shown below; identical to the corresponding sequence in the Swe1 genome). If an alternative nucleotide is supported by at least 5% of reads, the boxes are coloured proportionately, with the colours corresponding to those used for the different nucleotides in the genomic sequence shown below. Reads are represented in salmon or light blue, depending on their orientation. In those that do not match entirely the genomic sequence, non-consensus nucleotides are shown. For g7915.t1 (right), the gene is in the 3’-5’ orientation, while the alleles are given for the 5’-3’ sequence. (B) Summary of the number of reads supporting each allele in the 3 sets of reads.

**Supplementary Figure S5:**
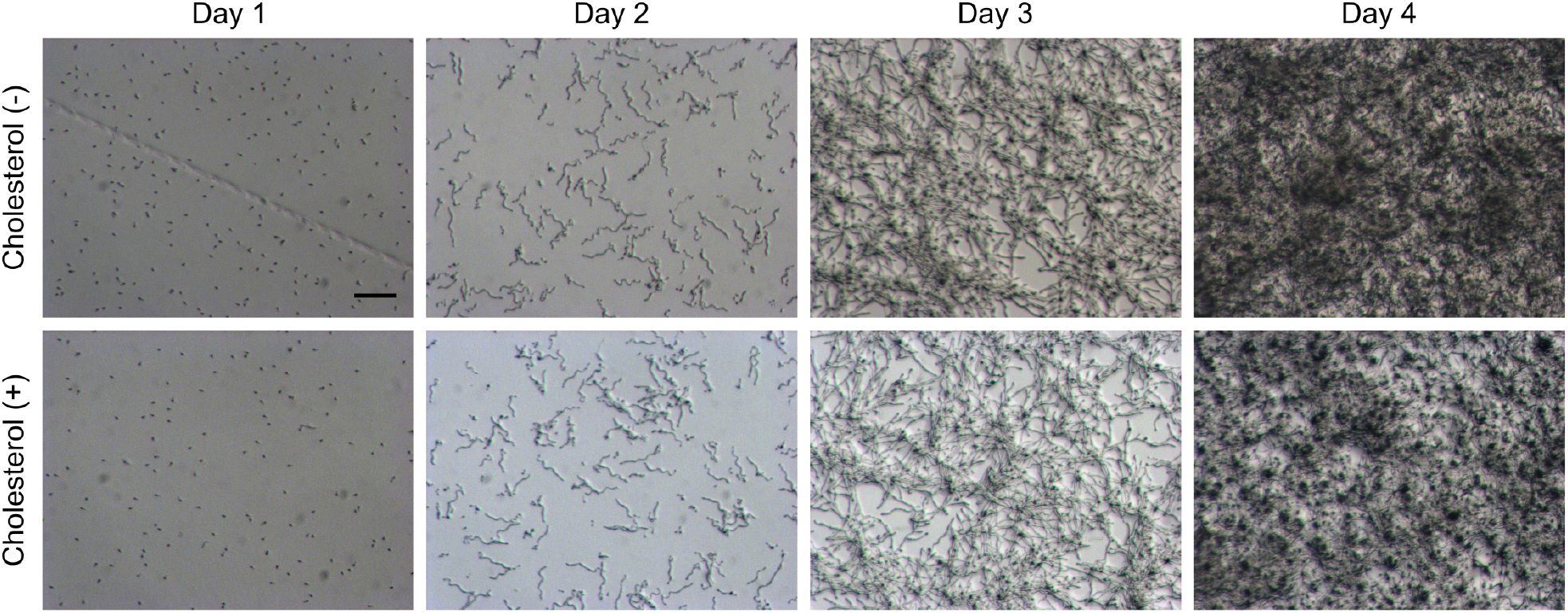
Growth of a *DcSre1* mutant is not altered by cholesterol supplementation. Swe3 carries a null allele of *DcSre1*. Fungal growth on standard NGM plates (+) or plates without (-) the normal 5 µg/l cholesterol supplementation was monitored at the indicated times. Scale = 100 µm.

**Supplementary Figure S6:**
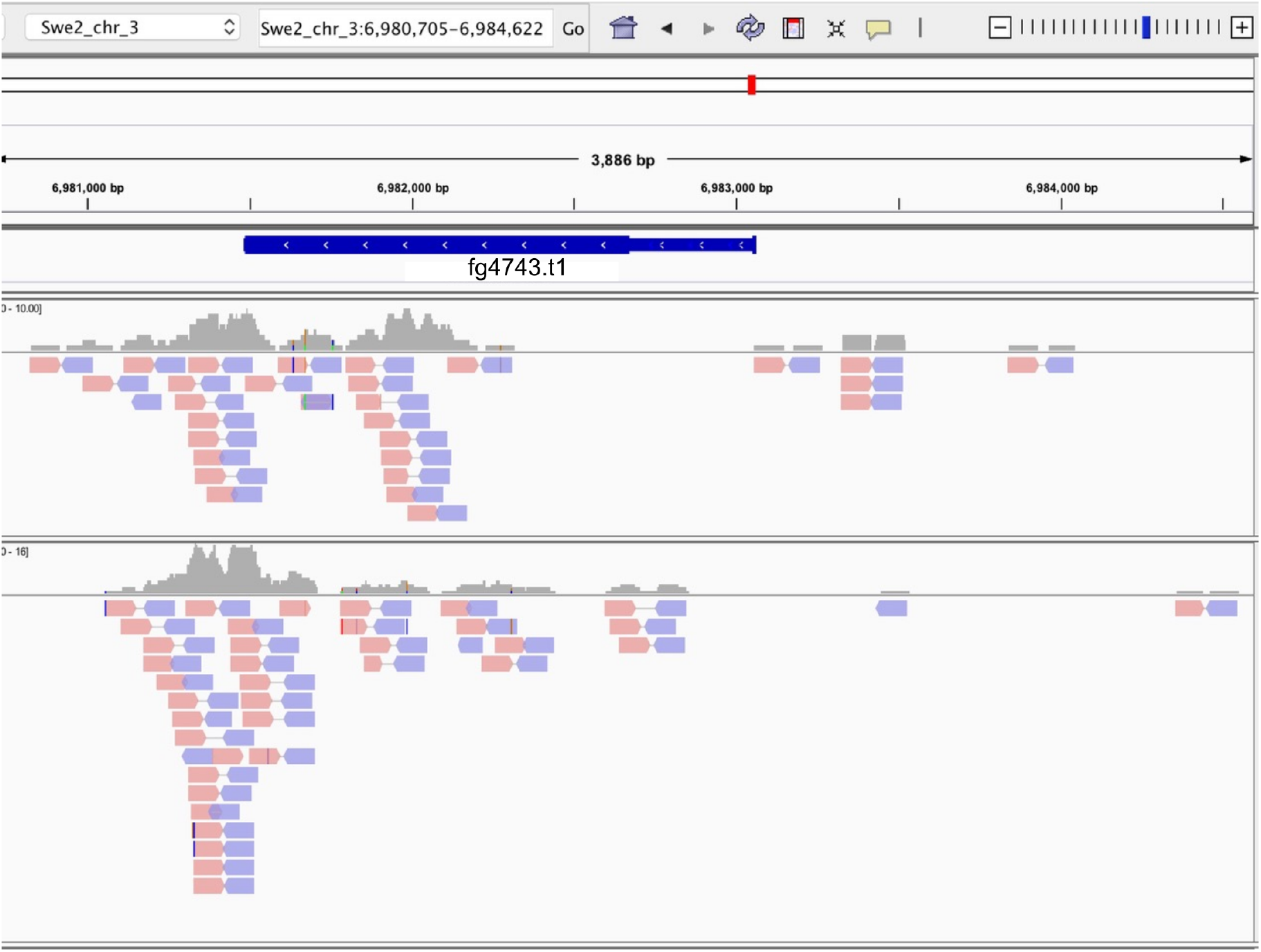
Imperfect RNAseq support for the current gene model of fg4743. Screen capture from IGV showing the mapping of two RNAseq read sets from mycelia (top) and spores (bottom) at the fg4743 (ODA79924.1) locus. The boxes at the top of each panel represent the read coverage at each nucleotide position. They are coloured grey if >95% of reads support the consensus sequence for the Swe2 genome. If an alternative nucleotide is supported by at least 5% of reads, the boxes are coloured proportionately, with the colours being the standard ones for the different nucleotides. Reads are represented in salmon or light blue, depending on their orientation, with coloured vertical lines indicating non-consensus nucleotides; paired reads are connected by a thin horizontal line.

**Supplementary Table 1:**
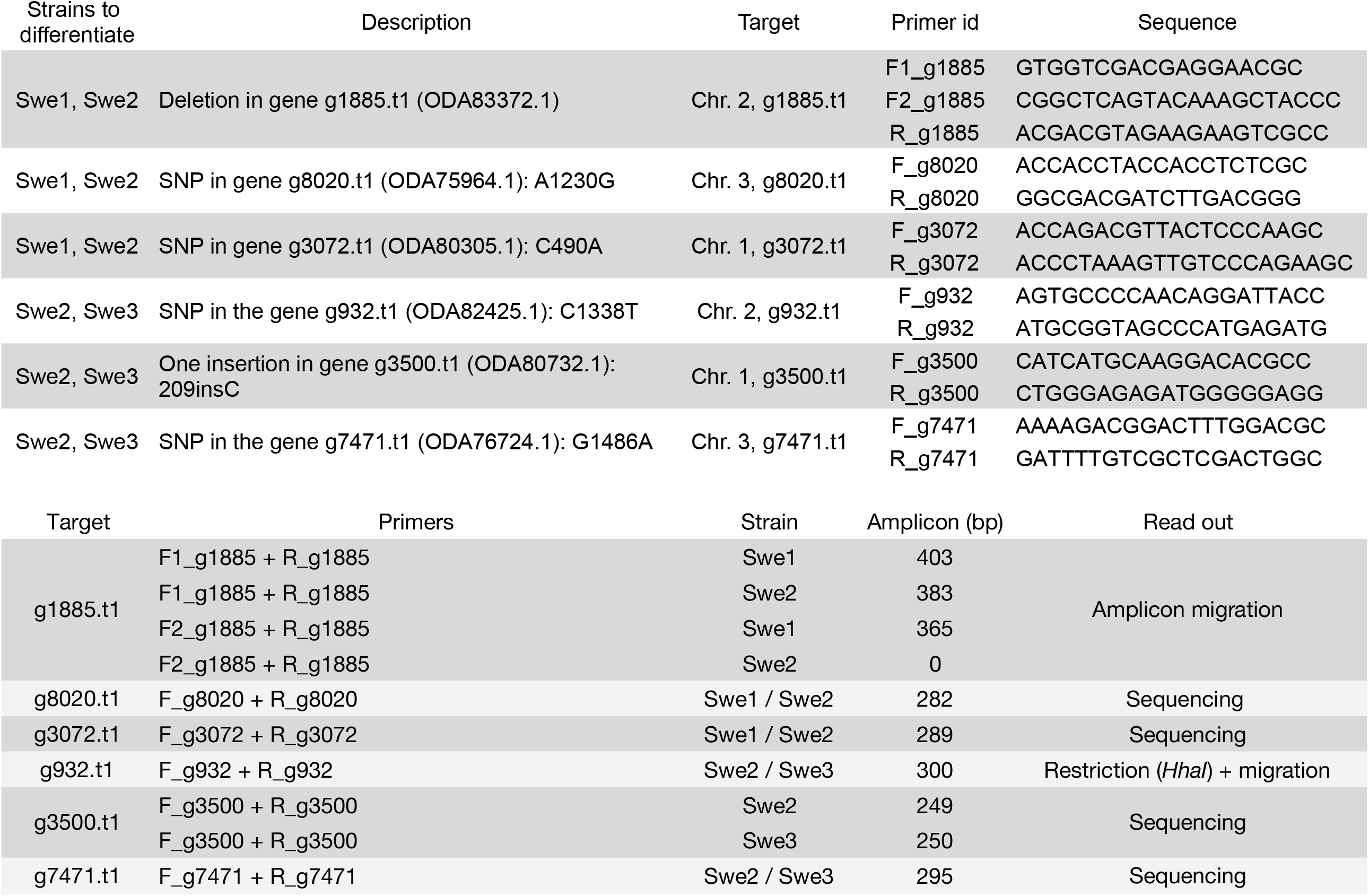
PCR tests used to genotype the *D. coniospora* Swe strains.

**Supplementary table 2:**
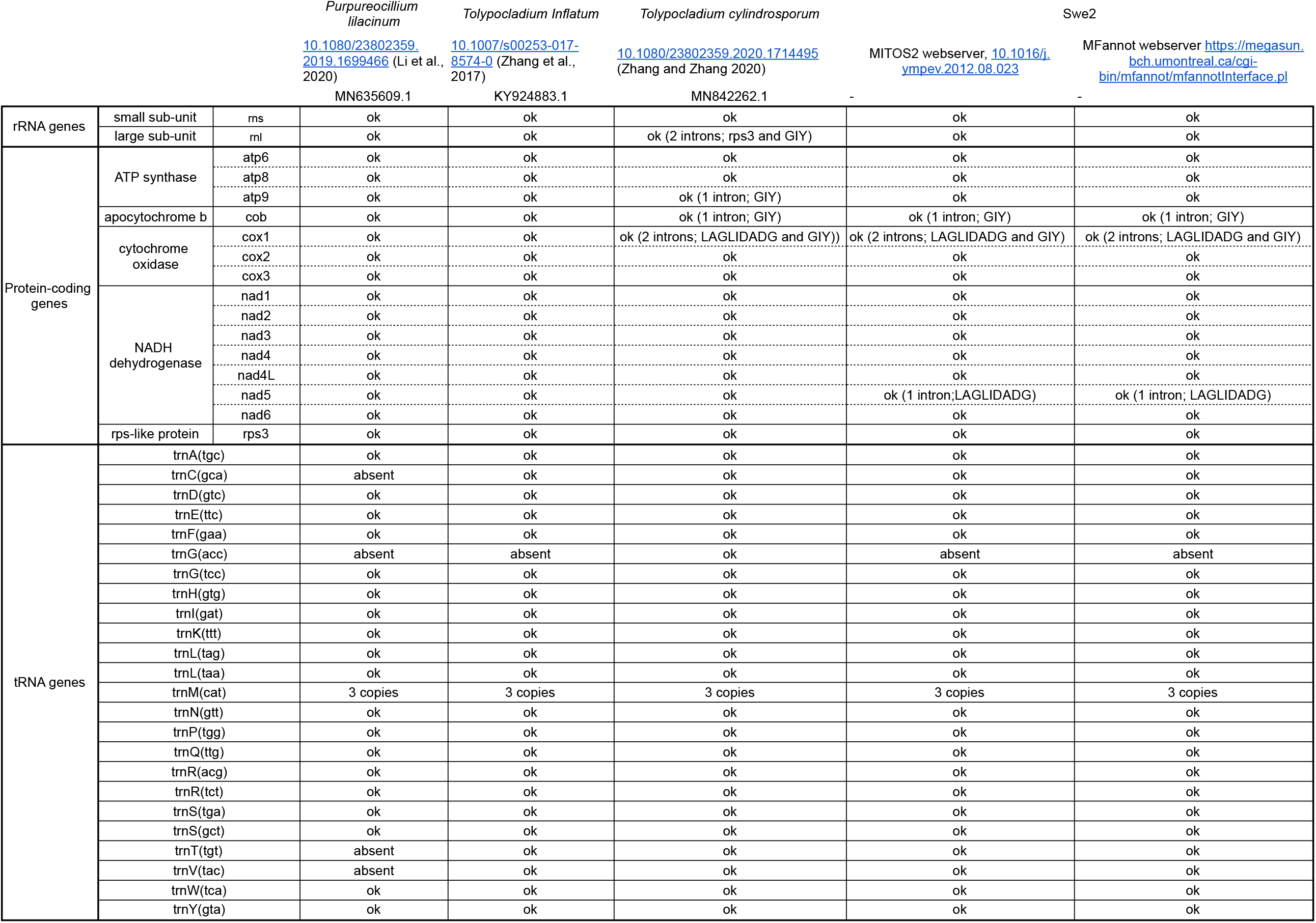
Presence/absence of mitochondrial genes found in Swe2 compared to 3 closely related fungi mitogenomes

**Supplementary table 3:** Status in the new assembly of Swe2 scaffolds that had not previously been incoprorated into the genome assembly

| Swe2 scaffold | length (nt) | presence of gene? | Insert in Swe2 | Length after trimming | remark |
| --- | --- | --- | --- | --- | --- |
| JYHR01000012.1 | 173 380 | yes | chromosome 3 | 160 167 |  |
| JYHR01000013.1 | 54 712 | yes | chromosome 1 | 50 350 |  |
| JYHR01000014.1 | 48 392 | yes | chromosome 3 | 43 976 |  |
| JYHR01000015.1 | 29 422 | yes | chromosome 1 | 28 782 |  |
| JYHR01000016.1 | 28 571 | yes | chromosome 3 | 28 571 |  |
| JYHR01000017.1 | 28 080 | yes | chromosome 3 | 21 992 |  |
| JYHR01000018.1 | 27 206 | yes | chromosome 3 | 27 206 |  |
| JYHR01000019.1 | 26 919 | yes | chromosome 3 | 26 919 |  |
| JYHR01000020.1 | 26 539 | yes | chromosome 2 | 26 539 |  |
| JYHR01000021.1 | 23 828 | yes | mitochondria | NA |  |
| JYHR01000022.1 | 23 058 | no | NA | NA |  |
| JYHR01000023.1 | 22 641 | yes | chromosome 3 | 22 641 |  |
| JYHR01000024.1 | 20 036 | yes | chromosome 3 | 16 817 |  |
| JYHR01000025.1 | 18 657 | yes | chromosome 3 | 18 657 |  |
| JYHR01000026.1 | 16 902 | yes | chromosome 3 | 16 902 |  |
| JYHR01000027.1 | 16 052 | yes | chromosome 3 | 16 052 |  |
| JYHR01000028.1 | 14 597 | no | NA | NA |  |
| JYHR01000029.1 | 14 514 | yes | chromosome 3 | 14 514 |  |
| JYHR01000030.1 | 14 170 | yes | chromosome 2 | 14 170 |  |
| JYHR01000031.1 | 14 066 | yes | chromosome 3 | 14 066 |  |
| JYHR01000032.1 | 9 307 | yes | chromosome 3 | 9 307 |  |
| JYHR01000033.1 | 9 128 | no | NA | NA |  |
| JYHR01000034.1 | 8 345 | no | NA | NA |  |
| JYHR01000035.1 | 8 253 | yes | chromosome 3 | 8 253 |  |
| JYHR01000036.1 | 6 725 | no | NA | NA |  |
| JYHR01000037.1 | 6 689 | no | NA | NA |  |
| JYHR01000038.1 | 6 421 | yes | chromosome 3 | 6 421 |  |
| JYHR01000039.1 | 6 407 | yes | NA | NA | Identical to JYHR01000038.1 |
| JYHR01000040.1 | 5 655 | no | NA | NA |  |
| JYHR01000041.1 | 5 653 | no | NA | NA |  |
| JYHR01000042.1 | 5 624 | no | NA | NA |  |
| JYHR01000043.1 | 5 249 | yes | NA | NA | Already present in the genome, assembly duplication |
| JYHR01000044.1 | 5 211 | yes | chromosome 3 | 5 211 |  |
| JYHR01000045.1 | 4 839 | no | NA | NA |  |
| JYHR01000046.1 | 3 936 | no | NA | NA |  |
| JYHR01000047.1 | 3 817 | yes | NA | NA | Already present in the genome, assembly duplication |
| JYHR01000048.1 | 3 440 | no | NA | NA |  |
| JYHR01000049.1 | 2 917 | no | NA | NA |  |
| JYHR01000050.1 | 2 571 | no | NA | NA |  |
| JYHR01000051.1 | 2 487 | yes | chromosome 3 | 2 487 |  |
| JYHR01000052.1 | 2 111 | no | NA | NA |  |
| JYHR01000053.1 | 2 040 | no | NA | NA |  |
| JYHR01000054.1 | 1 554 | no | NA | NA |  |
| JYHR01000055.1 | 1 460 | no | NA | NA |  |
| JYHR01000056.1 | 1 434 | no | NA | NA |  |
| JYHR01000057.1 | 1 343 | no | NA | NA |  |
| JYHR01000058.1 | 1 318 | no | NA | NA |  |
| JYHR01000059.1 | 1 285 | no | NA | NA |  |
| JYHR01000060.1 | 1 146 | no | NA | NA |  |
| JYHR01000061.1 | 1 041 | no | NA | NA |  |
| JYHR01000062.1 | 932 | no | NA | NA |  |
| JYHR01000063.1 | 902 | no | NA | NA |  |
| JYHR01000064.1 | 891 | no | NA | NA |  |
| JYHR01000065.1 | 825 | no | NA | NA |  |
| JYHR01000066.1 | 814 | no | NA | NA |  |
| JYHR01000067.1 | 812 | no | NA | NA |  |
| JYHR01000068.1 | 757 | no | NA | NA |  |
| JYHR01000069.1 | 722 | no | NA | NA |  |
| JYHR01000070.1 | 701 | no | NA | NA |  |
| JYHR01000071.1 | 636 | no | NA | NA |  |
| JYHR01000072.1 | 584 | no | NA | NA |  |
| JYHR01000073.1 | 556 | no | NA | NA |  |
| JYHR01000074.1 | 536 | no | NA | NA |  |
| JYHR01000075.1 | 505 | no | NA | NA |  |

Supplementary table 3 (continued): Swe2 assembly statistics
|  | Chromosomes | Unplaced scaffolds |
| --- | --- | --- |
| Genome size (Mb) | 31.75 | 0.128 |
| Sequences | 3 + MT | 38 |
| GC content (%) | 55.25 | 44.38 |
| % of unknowns (Ns) | 0.11 | 3.8 |
| No. predicted protein-coding genes | 8702 | 0 |

**Supplementary table 4:** ANI values for all pairs of Swe genomes from pyANI v0.29. Sequences were aligned with MUMmer (ANIm) or BLAST+ (ANIb). Tetranuccleotides correlations represent similarity with alignement-free method.

| Measure | Query strain | Swe1 | Swe2 | Swe3 |
| --- | --- | --- | --- | --- |
| ANIm | Swe1 | 100 | 99.9052 | 99.9896 |
|  | Swe2 | 99.9052 | 100 | 99.9060 |
|  | Swe3 | 99.9896 | 99.9060 | 100 |
| ANIB | Swe1 | 100 | 99.7694 | 99.9901 |
|  | Swe2 | 99.9155 | 100 | 99.9154 |
|  | Swe3 | 99.9820 | 99.7608 | 100 |
| Tetranucleotides correlations | Swe1 | 100 | 99.9709 | 99.9971 |
|  | Swe2 | 99.9709 | 100 | 99.9550 |
|  | Swe3 | 99.9971 | 99.9550 | 100 |

**Supplementary Table 5:** Results of BLAST searches of Dan2 sexual reproduction-related proteins

| Query | Query ID | Query size (aa) | Best BLASTP hit in Swe2 | Score | %ID | Alignment length (aa) | Query coverage (%) | Comment |
| --- | --- | --- | --- | --- | --- | --- | --- | --- |
| MAT1-1-1 | KYK59754.1 | 370 | - | - | - | - | - |  |
| MAT1-1-2 | KYK59755.1 | 353 | - | - | - | - | - |  |
| MAT1-1-3 | KYK59756.1 | 166 | ODA81607.1 | 71 | 40 | 75 | 45 | Alignment is restricted to the protein domain (HMG-box - PF00505) |
| RIP - rid1 | KYK59907.1 | 646 | ODA78704.1 | 776 | 91 | 416 | 64 |  |
| RIP - dim-2 | KYK56049.1 | 377 | ODA76538.1 | 598 | 86.4 | 397 | 100 |  |
| Late sexual development protein | KYK59150.1 | 355 | ODA77900.1 | 706 | 99.72 | 355 | 100 |  |
| Heterokaryon incompatibility | KYK58811.1 | 804 | ODA84177.1 | 1630 | 98.88 | 804 | 100 |  |
| Heterokaryon incompatibility | KYK56754.1 | 938 | ODA78419.1 | 1898 | 99.68 | 938 | 100 |  |
| Heterokaryon incompatibility | KYK56328.1 | 752 | ODA76785.1 | 1506 | 99.73 | 732 | 97 |  |
| Heterokaryon incompatibility - putative | KYK55209.1 | 749 | ODA82169.1 | 1546 | 99.73 | 749 | 100 |  |

| Protein of interest | Target | Best HSP (TBLASTN) | Best HSP localization | Best Query ID (UniProt ID) | Comment |
| --- | --- | --- | --- | --- | --- |
| MAT1-1-1 | Swe2 | no hit |  |  |  |
| MAT1-1-2 | Swe2 | no hit |  |  |  |
| MAT1-1-3 | Swe2 | ODA78559.1 | Chr. 1 | A7KPA5_COCPO | Query does not have the required protein domain (HMG-box - PF00505) |
| MAT1-2-1 | Swe2 | not hit | Chr. 1 | V9LW71_9HYPO |  |
| MAT1-2-2 | Swe2 | ODA81607.1 | Chr. 2 | A0A2K8C394_9PEZI | Alignment is restricted to the protein domain (HMG-box - PF00505) |
| MAT1-2-3 | Swe2 | not hit |  |  |  |
| MAT1-1-1 | Dan2 | KYK59754.1 | Chr. 1 (CM004174.1) | U3N6X4_TOLIN | Presence of the protein domain "Mating-type protein MAT alpha 1 HMG-box" (PF04769) |
| MAT1-1-2 | Dan2 | KYK59755.1 | Chr. 1 (CM004174.1) | Q1MVS6_TOLIN | Presence of the protein domain "Mating type protein 1-1-2" (PF17043) |
| MAT1-1-3 | Dan2 | KYK59757.1 | Chr. 1 (CM004174.1) | A7KPA5_COCPO | Query does not have the required protein domain (HMG-box - PF00505) |
|  |  | KYK59756.1 | Chr. 1 (CM004174.1) | Q1MVS7_TOLIN | Presence of of the protein domain "HMG-box" (PF00505) |
| MAT1-2-1 | Dan2 | KYK55799.1 | Chr. 3 (CM004176.1) | K4H8Z2_9HELO | Alignment is restricted to the protein domain (HMG-box - PF00505) |
| MAT1-2-2 | Dan2 | KYK55799.1 | Chr. 3 (CM004176.1) | A0A2K8C394_9PEZI | Alignment is restricted to the protein domain (HMG-box - PF00505) |
| MAT1-2-3 | Dan2 | no hit |  |  |  |

